# Inferring fungal growth rates from optical density data

**DOI:** 10.1101/2024.02.27.582053

**Authors:** Tara Hameed, Natasha Motsi, Elaine Bignell, Reiko J. Tanaka

## Abstract

Quantifying fungal growth underpins our ability to effectively treat severe fungal infections. Current methods quantify fungal growth rates from time-course morphology-specific data, such as hyphal length data. However, automated large-scale collection of such data lies beyond the scope of most clinical microbiology laboratories. In this paper, we propose a mathematical model of fungal growth to estimate morphology-specific growth rates from easy-to-collect, but indirect, optical density (OD_600_) data of *Aspergillus fumigatus* growth (filamentous fungus). Our method accounts for OD_600_ being an indirect measure by explicitly including the relationship between the indirect OD_600_ measurements and the calibrating true fungal growth in the model. Therefore, the method does not require *de novo* generation of calibration data. Our model outperformed reference models at fitting to and predicting OD_600_ growth curves and overcame observed discrepancies between morphology-specific rates inferred from OD_600_ versus directly measured data in reference models that did not include calibration.

**Author summary:** Quantifying fungal growth is essential for antifungal drug discovery and monitoring antifungal resistance. As fungal growth is complex, with fungal morphology (shape) dynamically changing over time, previous studies have quantified fungal growth by estimating growth rates during specific fungal morphologies (morphology-specific growth rates) or by mathematically modelling fungal growth. However, collecting time-series data that captures the morphological information required for mathematical model fitting or estimating morphology-specific growth rates is prohibitively time consuming for large-scale drug testing in most microbiology laboratories. Alternatively, fungal growth can be quickly, although indirectly, quantified by measuring the optical density (OD) of a broth culture. However, changes in OD are not always reflective of true changes in fungal growth because OD is an indirect measure. This paper proposes a method to model fungal growth and estimate a morphology-specific growth rate from indirect OD_600_ measurements of the major mould pathogen, *Aspergillus fumigatus*. We explicitly model the relationship between measured indirect OD_600_ data and true fungal growth (calibration). The presented work serves as the much-needed foundation for estimating and comparing morphology-specific fungal growth rates in varying antifungal drug concentrations using only OD_600_ data.

## Introduction

Fungal infections affect over a billion people worldwide [1] and remain a global threat to human health [2]. Combatting fungal infections is essential for treating patients with life-threatening invasive infections [1]. In the clinic, the invasive fungal infection, invasive aspergillosis, is widely prevalent [1] and associated with unacceptably high mortality rates [2] of around 29% [3], even with treatment. Without appropriate treatment, invasive aspergillosis can result in 100% mortality, as seen with COVID-associated-pulmonary-aspergillosis [4]. Whilst most fungal infections are treated with antifungal drugs [5,6], the issue of antifungal resistance [7], spanning to multi-drug resistance, may shrink an already small pool of treatment options for patients [8]. To avoid patients being left with no treatment options, there is a real need to locate new antifungal drugs and for increased antifungal stewardship [7].

Antifungal stewardship requires detecting and quantifying fungal growth in different conditions to evaluate potential resistance or drug efficacy. Multiple fungal pathogens have recently been identified as critical threats to public health by the world health organisation (WHO) [9]. These fungi can exhibit complex growth dynamics, such as filamentation, due to morphogenesis [10,11], where morphogenesis is also promoted by exposure to antifungal drugs [12]. The current standards for antifungal susceptibility testing for filamentous fungi, set by the Clinical and Laboratory Standards Institute (CLSI) and the European Committee for Antimicrobial Susceptibility Testing (EUCAST), involve evaluating minimum inhibitory/fungicidal/effective concentrations (MIC/MFC/MEC) *in vitro* [13–15], which is the minimum drug concentration required to reduce or alter the morphology of fungal biomass within a certain time [15–18]. However, further development of antifungal susceptibility testing for filamentous fungi is required for the following two reasons. First, the current CLSI and EUCAST standards include subjective measurements that depend upon visual inspection of changes in growth [19]. Second, the current guidelines cannot capture time-dependent antifungal activity or antifungal activity specific to a particular fungal morphology (morphology-specific antifungal activity). The MIC/MFC/MEC do not describe temporal dynamics of fungal growth because they are based upon a single end-point measurement. These metrics also cannot capture morphology-specific antifungal activity while fungal morphology evolves over time. Popular antifungal drugs, such as azoles, have morphology-specific mechanisms of action [20] and different fungal morphologies have been noted to be future antifungal targets [21]. We need a more quantitative, temporal, and morphology-specific evaluation of fungal growth to further mechanistic understanding of both differential growth of filamentous fungi and a drug’s antifungal activity.

Several previous studies have calculated fungal growth rates to quantitatively evaluate temporal changes in fungal growth in different experimental conditions [22–25]. Growth rates specific to a fungal morphology (morphology-specific growth rates), such as germination rates of fungal spores or proliferation rates of hyphae, can be estimated from morphology-specific growth data to evaluate morphology-specific changes in fungal growth over time. For example, hyphal extension rates have been derived from hyphal microscopy data to investigate hyphae-specific growth defects in mutant strains of *Aspergillus fumigatus* [22]. Such morphology-specific growth rates can also be estimated from mechanistic models of fungal growth fit to directly measured time-course data. For example, hyphal growth rates have been estimated using a fungal growth model fit to hyphal area image data [26]. Mechanistic models that include morphology-specific growth mechanisms [23,26,27] can not only estimate morphology-specific growth rates, but also further understanding of morphological dynamics during growth [26]. However, to the best of our knowledge, estimating morphology-specific growth rates and developing mechanistic models of dynamic filamentous fungal growth are not routine in clinical microbiology laboratories potentially because it is too difficult to quickly collect the required time-course data that encapsulates morphological information or directly measures dynamic fungal growth.

The acquisition and analysis of directly measured time-course morphology data requires specialist technology and analyses that lie beyond the scope of clinical microbiology laboratories. To collect morphology time-course data, fungal growth is directly measured over time using techniques such as enumerating colony forming units (CFUs), microscopy, or histology. These direct data collection methods are only feasible for large-scale experimentation in specialist research microbiology laboratories because of the high cost of technologies and skill-level required to perform automated direct measurement of fungal growth. Technological developments have been made to aide processing of fungal image data [25], such as FIJI [28] or fungal feature tracker (FFT) [29]. But they are either not fully automated, relying on point-and-click methods, or are still not used in clinical microbiology labs due to difficulty of use or implementation.

Instead of directly measuring fungal growth to collect time-course fungal growth data, indirect measures of fungal growth can be obtained for growth rate estimation or mechanistic modelling by measuring optical density (OD). Collecting OD data is an automated, quick, and popular method to obtain time-dense fungal growth data that is widely used for estimating microbial growth rates [30] and is amenable to high-throughput phenotyping and antimicrobial susceptibility studies [31]. For fungi, the use of OD data to directly and empirically infer fungal growth rates is commonplace. Moreover, simple mechanistic models of fungal growth, such as logistic [32] and Gompertz [33] population growth models, have been directly fit to OD data to estimate fungal growth rates [23]. These simple mechanistic models could be extended to include morphology-specific growth mechanisms to estimate morphology-specific growth rates, as has been done with mechanistic models fit to directly measured data [26]. However, all these approaches to estimate growth rates from OD, both empirically or using modelling, harbour intrinsic inaccuracies because OD is an indirect measure of growth whereby changes in OD measurements may not even be reflective of true fungal growth and changes in fungal morphology are generally overlooked [11]. To collect data that has a higher fidelity to true fungal growth, OD data can be converted to a direct measure of growth using OD values at known fungal concentrations [34,35] (calibration). However, it is difficult for clinical microbiology laboratories to routinely collect time-dense direct measures of fungal growth required for robust calibration because automated collection of such data is not yet standard.

In this paper, we aim to model filamentous fungal growth and estimate morphology-specific fungal growth rates from OD measured at a wavelength of 600nm (OD_600_) without collecting calibration data but still taking into consideration that OD is an indirect measure of growth. We proposed a model of fungal growth that treats measured OD_600_ as an observed variable distributed around a linear transform of the true unobserved fungal concentration, thereby explicitly including calibration in the model structure. We demonstrate the model’s ability to fit to and predict OD_600_ using time-course OD_600_ data for the filamentous fungus, *A. fumigatus,* and evaluated the model’s estimated morphology-specific growth rate using directly measured data of *A. fumigatus* growth.

## Results

### OD_600_ data and directly measured fungal data

We obtained time-course OD_600_ data of fungal growth from an *in vitro* experiment of five initial inocula of uninucleate *A. fumigatus* spores (0, 2 × 10^2^, 2 × 10^3^, 2 × 10^4^ and 2 × 10^5^ spores) suspended in 200μl of fungal (f) Roswell Park Memorial Institute (RPMI) media (0, 1, 10^1^, 10^2^ and 10^3^ [nuclei (N)/μl]) (**Fig 1a**). We also collected two sets of directly measured *A. fumigatus* growth time-course data: the hyphal length (HL, **Fig 1b**) and nuclear count (NC, **Fig 1c**). HLs (μm) were measured using the software ImageJ [36] from microscopy images of wells using an initial inoculum of 1 [N/μl]. NC data was obtained by imaging a fluorescent strain of *A. fumigatus* with an initial inoculum of 10^1^ [N/μl] and counting the nuclei per hypha using the particle counter method in ImageJ [36].

**Fig 1:**
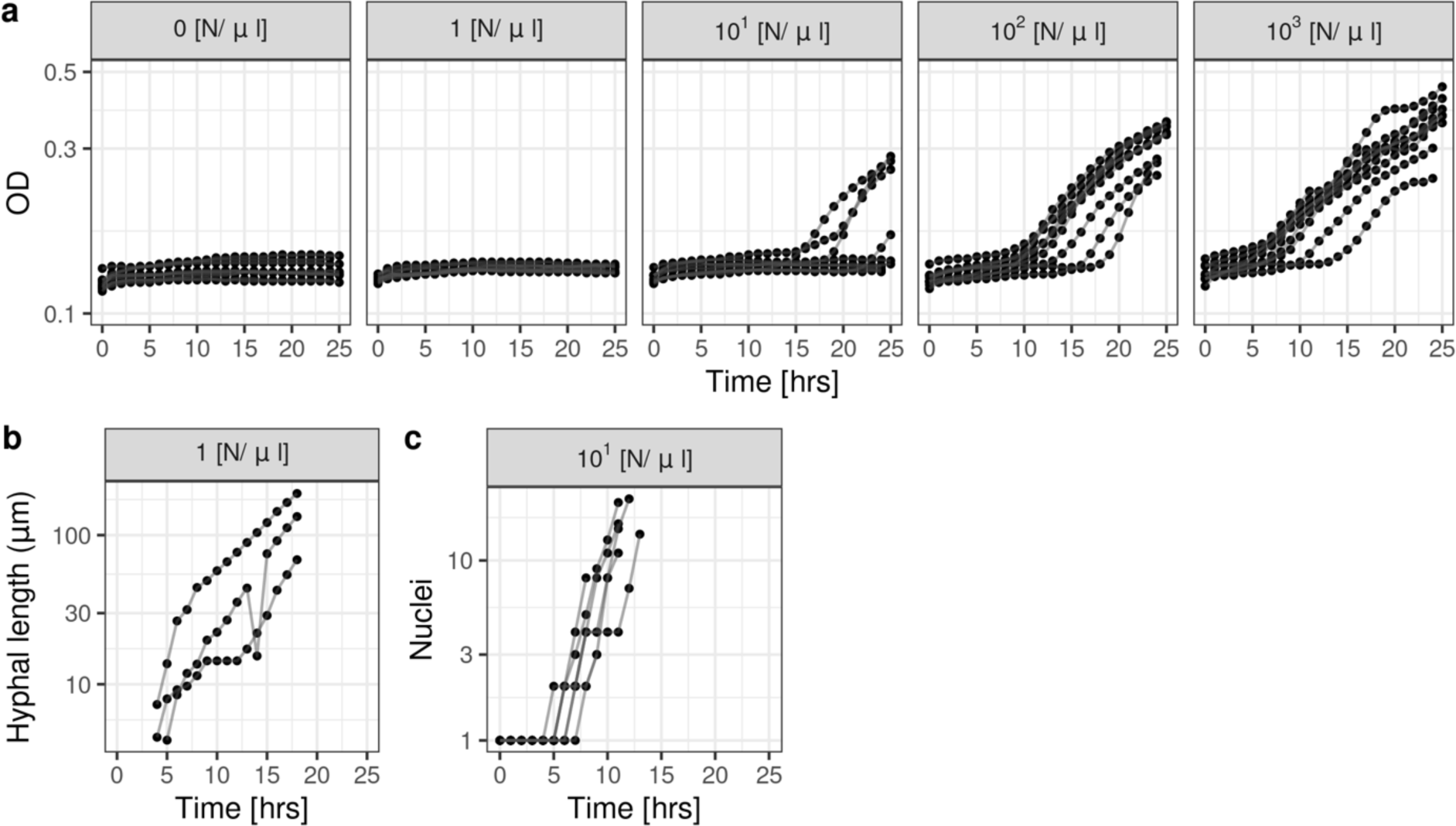
OD_600_ and directly measured fungal growth data. **(a)** *A. fumigatus* OD_600_ data (dots) recorded for 25 hours with initial inoculums of 0, 1, 10^1^, 10^2^, 10^3^ [N/μl] (left to right). (**b**) Hyphal length data of a single *A. fumigatus* hypha from wells with an initial inocula of 1 [N/μl]. (**c**) Nuclear count data of a single H1 *A. fumigatus* (fluorescent strain) hypha from wells initiliased with 10^1^ [N/μl]. Data from a single replicate is joined by a line. The y-axes are in the log_10_ scale.

The growth curves of the OD_600_ data differ from those of the HL and NC data. The OD_600_ data demonstrated little-to-no growth in the first 15 hours for lower fungal inocula (1-10^1^ [N/μl]), with values similar to those for 0 [N/μl] (fRPMI only control), and did not increase as rapidly as for a higher fungal inoculum (10^3^ [N/μl]). However, the HL and NC data demonstrated an increase in fungal growth in wells initialised with lower inocula (1-10^1^ [N/μl]) at as early as 5 hours. For the higher fungal inoculum of the OD_600_ data (10^3^ [N/μl]), the data demonstrated qualitatively similar dynamics to all of the fungal inocula in the directly measured data (HL, NC), with fungal growth rapidly increasing from around 5 hours. The different dynamics observed in OD_600_ data for the different fungal inocula are attributed to OD_600_ being an indirect measure of fungal growth rather than reflective of true fungal growth. The OD reader does not have the sensitivity required to detect true fungal growth in the wells initialised with lower fungal inocula for longer time-periods than wells initialised with the higher inocula.

### A logistic model fit to OD_600_ resulted in model misfit and growth rates distinct from directly measured data

We first outline the limitations of not considering OD being an indirect measure of growth when fitting fungal growth models to OD data. We assessed the fit of a simple population growth model (Logistic-OD model) to the OD_600_ data. The Logistic-OD model treated the measured OD_600_ as being distributed around the true fungal concentration in the wells [N/μl], which was modelled using a logistic function (see Methods). The logistic function was modified to model morphology-specific fungal growth by beginning growth after τ [hours], which reflects dormant spores germinating into growing hyphae τ [hours] after wells are inoculated with spores (referred to as a “delay” hereafter). Hence, the logistic function includes the growth rate during hyphal growth (hyphal growth rate) as a parameter to be estimated.

The Logistic-OD model was unable to fit to the measured OD_600_ data well (**Fig 2a**). The posterior predictive distribution (median and 80% credible intervals (CIs)) of the fitted logistic model did not include any of the data from a low-density fungal culture (1 [N/μl]) after 20 hours, where OD_600_ measurements registered little-to-no growth (**Fig 2a**). This was also observed when we fit alternative fungal growth models (an exponential model with and without a delay) to the OD_600_ data (**Fig S1**). The model misfit was unsurprising because the model cannot account for OD_600_ dynamics dependent on inoculum size. The initial value of the logistic function could not depend on the initial fungal inocula in the well because there was no observed difference in OD_600_ at *t* = 0 hours. Additionally, we assume that none of the growth parameters in the model (hyphal growth rate (β) or the delay (τ)) depend on the initial inoculum size because there is no clear evidence to suggest this in our HL and NC data or in the literature. We found only one study with data showing a correlation between inoculum size and *A. fumigatus* germination rates [37], to the best of our knowledge. Fitting the Logistic-OD model only to the high inoculum data (10^3^ [N/μl]) removes the need for the model to account for inoculum size-dependent OD_600_ dynamics, and the model could fit to the data well. The posterior predictive distribution of the Logistic-OD model included more of the high inoculum data when the model was fit to only the high inoculum data (**Fig 2b**), as opposed to all the data.

**Fig 2:**
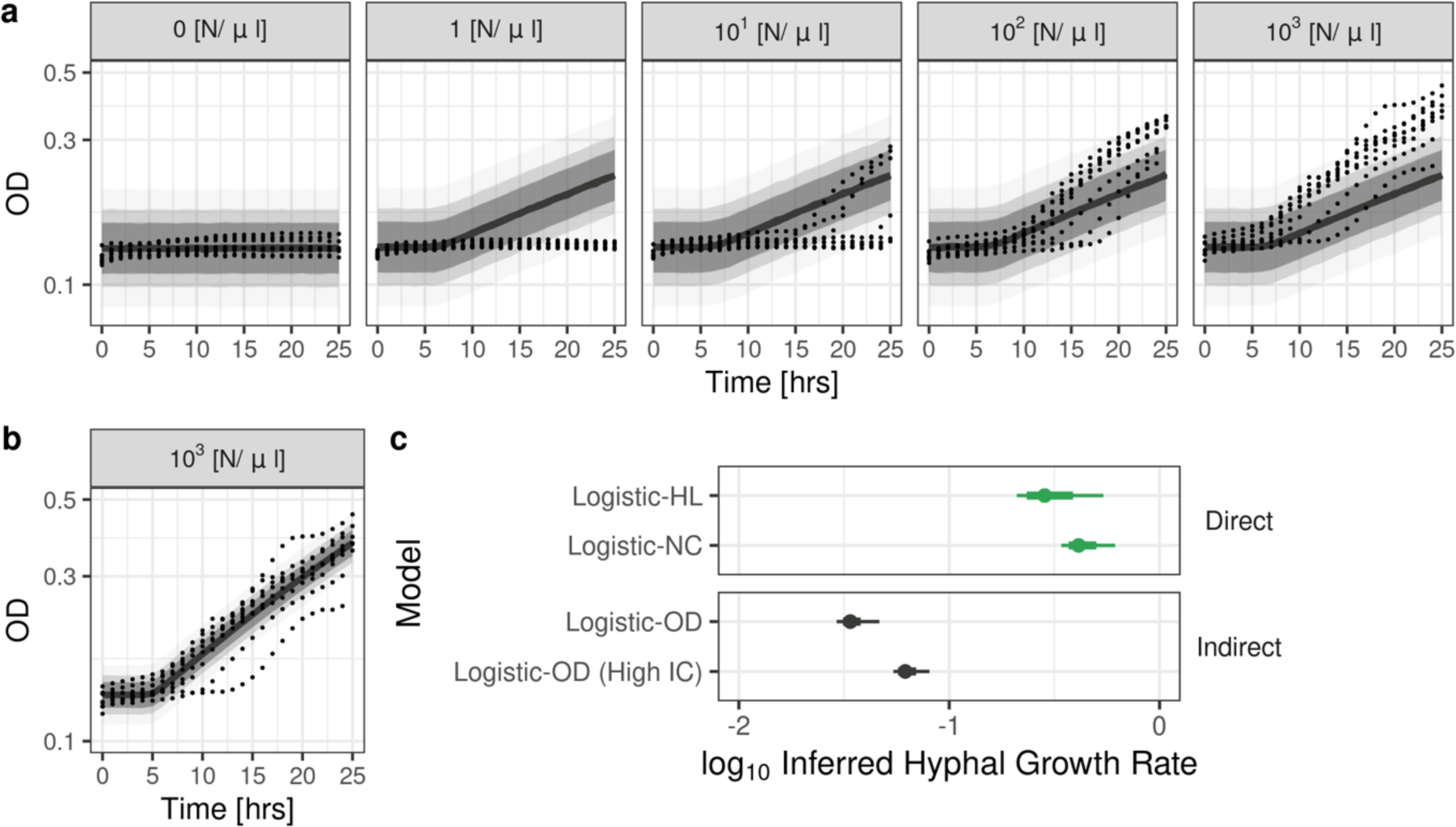
Logistic-OD model fit to OD_600_ data and estimated growth rates. Posterior predictive distribution of the Logistic-OD model fit to the OD_600_ data (black dots) with **(a)** all the initial fungal inoculums (0, 1, 10^1^, 10^2^, 10^3^ [N/μl], left to right) and (**b**) only the high fungal inoculum (10^3^ [N/μl]). Solid lines and shades are medians and 95%, 80%, 60% credible intervals, respectively. The y-axes are in the log_10_ scale. (**c**) Hyphal growth rates (reference rates, green) estimated using a logistic model fit to direct data (Logistic-HL fit to hyphal length and Logistic-NC fit to nuclear count data, respectively) and those inferred from the Logistic-OD model fit to the indirect OD_600_ data, either all data (Logistic-OD) or only the data with high fungal inocula (Logistic-OD (High initial condition (IC))). Medians (dot), 95% (thin line) and 80% (thick line) credible intervals are shown. The hyphal growth rates of the Logistic-OD model are distinctly lower than the reference growth rates. All the inferred growth rates are shown on log_10_ scale.

We next evaluated the ability of the Logistic-OD model to estimate a morphology-specific growth rate (hyphal growth rate). We estimated hyphal growth rates from two logistic models fit to directly measured data from two experiments, namely the Logistic-HL model fit to time-course microscopy-based HL data and the Logistic-NC model fit to NC data (**Fig 2c**, Direct). We refer to those rates as reference rates. The reference rates’ 95% and 80% CIs overlapped, and their medians are in the same order of magnitude (**Fig 2c**, Direct). However, the hyphal growth rates inferred from the Logistic-OD model, either fit to all the OD_600_ data or only the high fungal inoculum (10^3^[N/μl]) data, had their median and quantiles being distinctly lower and outside of the 95% CIs of the reference rates (**Fig 2c**, Indirect).

Overall, fitting a simple population growth model that does not account for OD being an indirect measure to uncalibrated OD_600_ data resulted in both model misfit and a distinctly lower growth rate compared to those estimated from direct measures of fungal growth (HL and NC).

### Logistic-OD-calibration model that incorporates calibration successfully fits to OD_600_ data

To combat the model misfit observed with the Logistic-OD model, we proposed a fungal growth model (Logistic-OD-calibration model) that considers the OD_600_ data being an indirect measure of growth (see Methods). The Logistic-OD-calibration model incorporates calibration of OD by explicitly modelling the measured OD_600_ as an observed variable distributed around a linear transform of the true fungal concentration [N/μl]. The linear transform is described with two parameters, a background correction, B, and a proportionality constant, 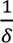. The model uses the same logistic function as the Logistic-OD model to describe the true fungal growth, which had an explicit hyphal growth rate and a time delay of τ [hours] to reflect spore germination. In this way, we explicitly modelled the relationship between OD_600_ and fungal growth (calibration curve), whilst retaining a biologically interpretable growth rate in the model.

The Logistic-OD-calibration model was able to successfully fit to all of the time course OD_600_ data we collected, as confirmed by posterior predictive checks (**Fig 3**). The model’s posterior predictive distribution captured flat or delayed OD_600_ growth curves for lower fungal inocula (0-10^1^ [N/μl]) and sigmoidal growth curves for a higher inoculum (10^3^ [N/μl]), whilst assuming the same underlying latent logistic function for fungal growth in each well (**Fig 3**).

**Fig 3:**
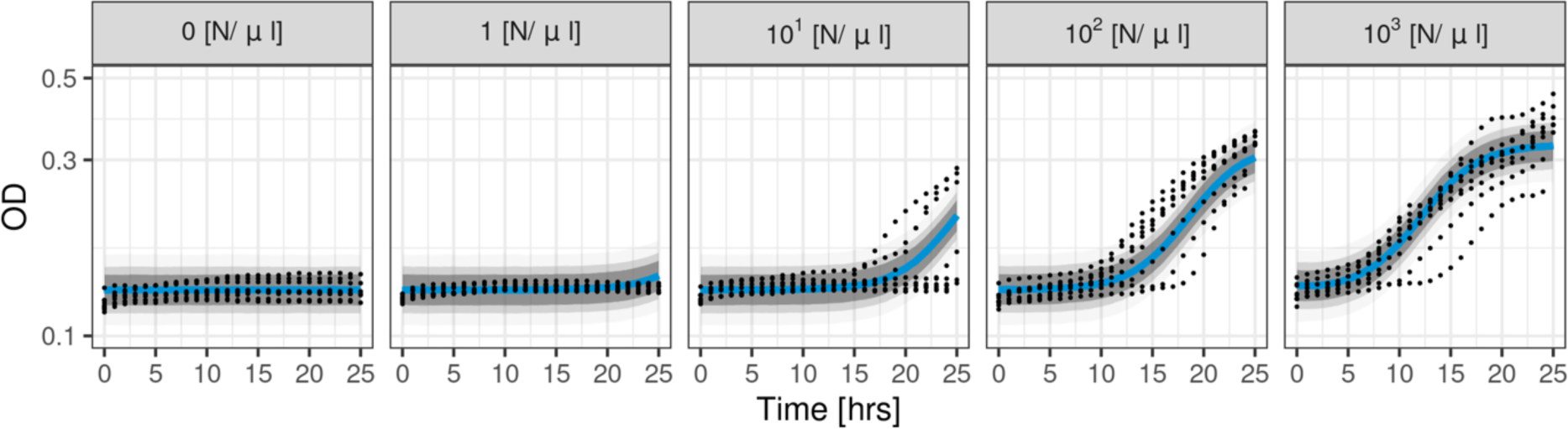
Logistic-OD-calibration model fit to OD_600_ data. Posterior predictive distribution is shown with medians (solid blue lines) and credible intervals of 95%, 80%, 60% (shaded in grey). Black dots are the observed OD_600_ data from wells initialised with different fungal inocula (0, 1, 10^1^, 10^2^, 10^3^ [N/μl], from left to right). The y-axis is in the log_10_ scale.

### Modelling calibration improved model fit and predictions

We then quantitatively assessed the benefit of including calibration in our model in terms of the model fit to the OD_600_ data and the prediction of the OD_600_ values. We quantified model fit by root mean squared error (RMSE) on training data and model prediction by relative log predictive density (LPD) and RMSE on entire replicates of time-course OD_600_ held-out during cross validation (CV). The lower the RMSE and relative LPD are, the better performing the model is.

Our Logistic-OD-calibration model demonstrated markedly lower RMSE and relative LPD, compared to the Logistic-OD model that did not include calibration (**Fig 4**, reference logistic model without calibration), confirming the benefit of including calibration in the model. Our model also demonstrated lower RMSE and relative LPD compared to other reference models that did not incorporate calibration and were directly fit to OD_600_, including a Gompertz and Gaussian Process (GP) model that are popular parametric and non-parametric models used in OD modelling literature [34,38–41], respectively (**Fig 4**, reference models without calibration). When we used OD_600_ data only from wells inoculated with the highest fungal inoculum (10^3^ [N/μl]), models that did not include calibration (logistic, Gompertz and GP) achieved comparable RMSE and relative LPD values with our model (**Fig S2**). These results suggest that including calibration in our model is beneficial when the data is likely affected by how the OD reader measures fungal growth, for example fungal growth falling below a detection limit, but it does not worsen model performance when the data was not obviously affected by the OD reader.

**Fig 4:**
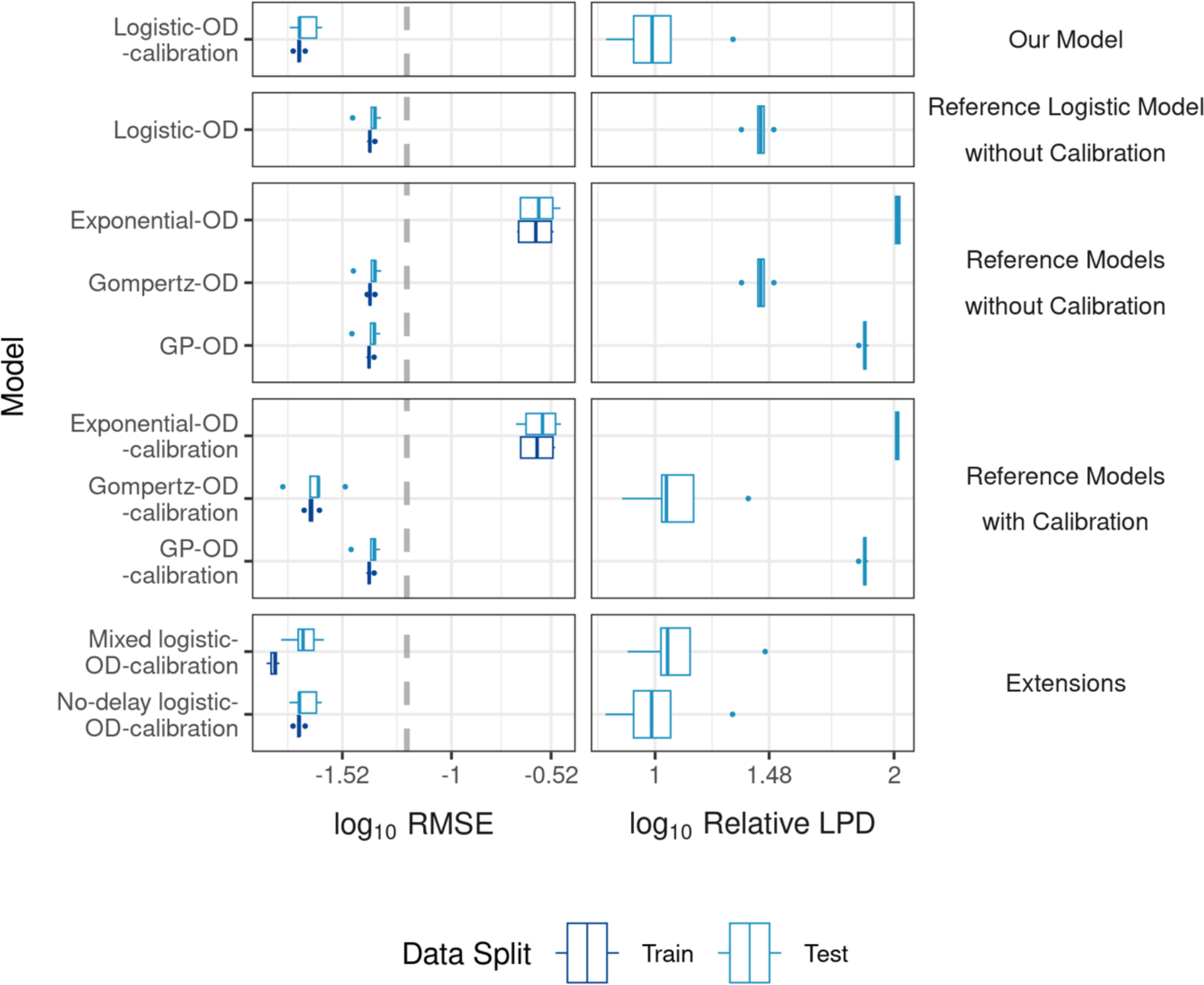
Model fit and prediction. Model fit and prediction are quantified by RMSE (three significant figures (3.s.f.)) averaged over training (dark blue) and testing (light blue) OD_600_ replicates, respectively, for each fold in 5-fold cross validation stratified by replicates (left). The dotted grey line indicates the RMSE of an intercept model. Model prediction is quantified by Relative LPD ((maximum LPD) - LPD) (3.s.f.) with LPD averaged over test replicates (right) for each fold. Lower values indicate better fitting and predictive performance for both metrics.

We next verified the benefit of our choice of a logistic function for the true fungal growth by comparing our model to reference models that included calibration but with different growth models for the fungal growth, including an exponential function, Gompertz model and a GP (**Fig 4**, reference models with calibration). Our model achieved better predictive performance, shown by lower relative LPD, than the exponential, Gompertz and GP models, achieving higher predictive power (relative LPD) than all the reference models. The GP-OD and GP-OD-calibration models, and similarly the Logistic-OD-calibration and No-delay logistic-OD-calibration models, achieve different RMSE and relative LPD (**Table S1**) but are visually indistinguishable (**Fig 4**) due to similar model fits to the OD_600_ data during CV (**Fig S3**).

Introducing random effects on the background OD_600_ data as measured from control wells containing only fRPMI (modelling a background media value per replicate instead of estimating the mean background level of the replicates) worsened the model performance (relative LPD), while removing a growth delay (*τ* = 0) did not result in much difference in model performance (**Fig 4**, extensions). We therefore decided to use the Logistic-OD-calibration model with a growth delay to infer a fungal growth rate because *A. fumigatus* is known to exhibit a period of no growth [42], which can be seen in the HL and NC data. Including the delay allows us to differentiate dormant fungal spores (*t* ≤ *τ*) from growing hyphae (*t* > *τ*) and, hence interpret the growth rate as a hyphal growth rate, which has a morphological and biological interpretation.

### The logistic-OD-calibration model could overcome the discrepancy between growth rates estimated from OD_600_ and directly measured data

Given that our Logistic-OD-calibration model could fit to all the collected OD_600_ data and could predict held-out OD_600_ data during CV better than the reference models, we next evaluated the hyphal growth rate inferred by our model from OD_600_ data, by comparing our inferred (Logistic-OD-calibration) hyphal growth rate to the reference growth rates estimated from logistic models fit to HL and NC data (Logistic-HL and Logistic-NC) (**Fig 5a**). The estimated median and quantiles for Logistic-OD-calibration model’s hyphal growth rate overlapped with the CIs of the reference growth rates. This is in contrast with a distinctly lower median rate estimated by the Logistic-OD model without calibration, suggesting that the calibration included in our model was crucial to overcome the discrepancy between the Logistic-OD model’s growth rate and the reference rates.

**Fig 5:**
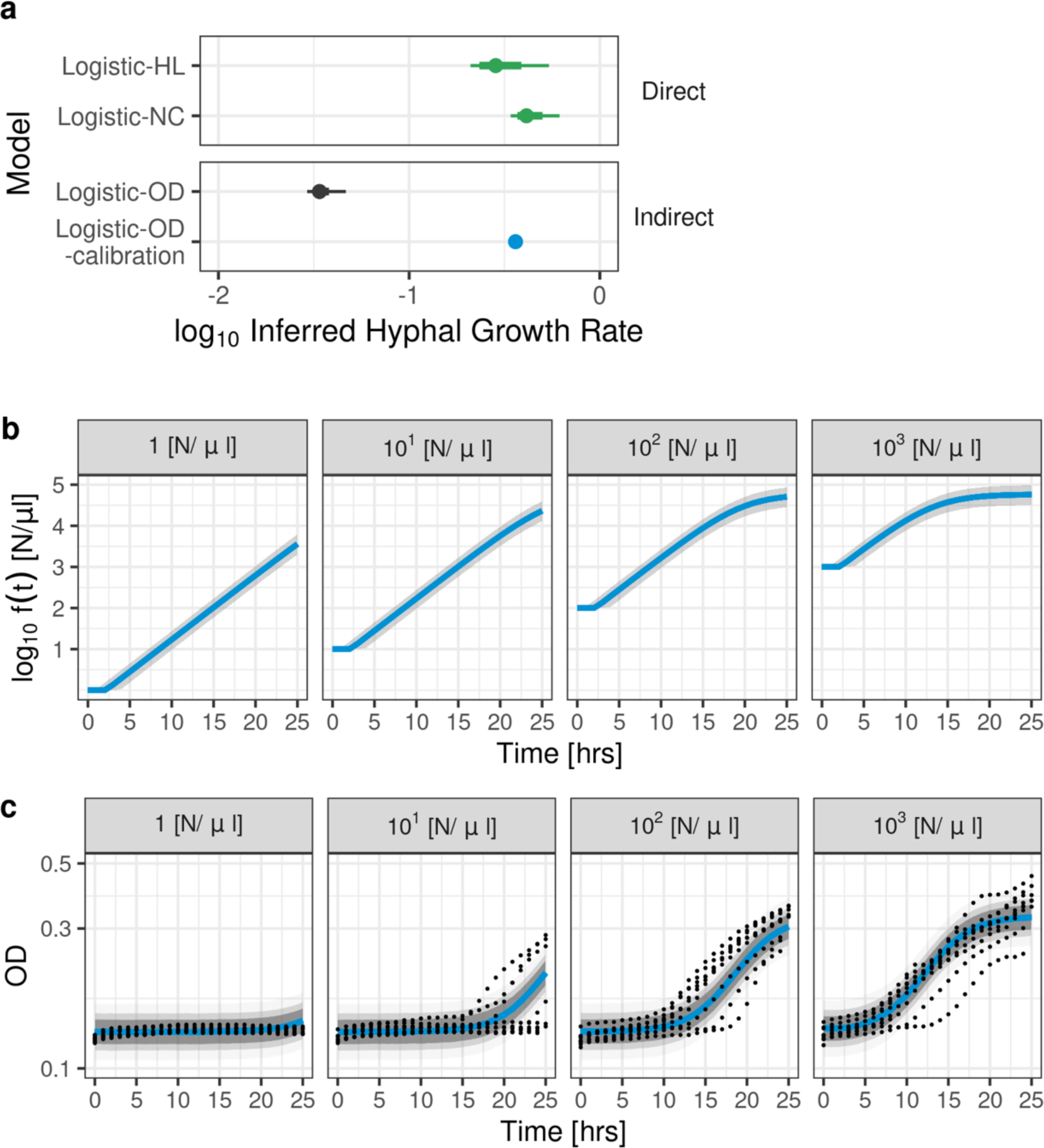
Hyphal growth rate estimated by Logistic-OD-calibration model and inferred true fungal growth from model fit to OD_600_. **(a)** Medians (dot), 95% (thin line) and 80% (thick line) credible intervals of hyphal growth rates (reference rates, green) estimated from logistic models fit to direct data (Logistic-HL and Logistic-NC fit to hyphal length and nuclear count data, respectively) compared to the estimated hyphal growth rate from our Logistic-OD-calibration model fit to all the OD_600_ data (blue). The 80% and 95% credible intervals ([0.352, 0.371] and [0.348, 0.376] 3.s.f., respectively) of the growth rate inferred using our Logistic-OD-calibration model (blue) overlap with the credible intervals of reference growth rates (green), but the credible intervals of the growth rate of the Logistic-OD model (black) do not. (**b**) Inferred true fungal growth, *f*(*t*), in [N/μl] (log_10_ scale) from our Logistic-OD-calibration model when modelling fungal growth from OD_600_ data measured from wells inoculated with 1, 10^1^, 10^2^, 10^3^ [N/μl] (left to right). Solid lines and shades are medians and 95% credible intervals, respectively. (**c**) Logistic-OD-calibration model fit (posterior predictive distribution, medians shown in solid blue line and 95%, 80%, 60% credible intervals shaded) to measured OD_600_ (dots) from wells inoculated with 1, 10^1^, 10^2^, 10^3^ [N/μl] (left to right), where the OD_600_ data was treated as distributed around a linear transform of the true fungal growth, *f*(*t*) (shown in (**b**)). The y-axis is in the log_10_ scale.

For OD_600_ data from the high fungal inoculum, the posterior predictive distribution of the Logistic-OD model without calibration successfully captured the data (**Fig 2b**) but the estimated growth rates were further from the reference rates compared to the rates estimated by our model with calibration (**Fig S4**). The Gompertz-OD model fit to all the OD_600_ data fails to infer similar initial hyphal growth rates to the reference hyphal growth rates estimated from the HL and NC data (**Fig S5**). When the model was fit to only the high initial inoculum OD_600_ data, the estimated initial growth rate had overlapping 95% CIs with the reference growth rates (**Fig S5**). However, if the calibration was modelled (Gompertz-OD-calibration model), the initial growth rates inferred from both all the OD_600_ data and only the high initial inoculum had overlapping 80% CIs with both the reference rates, and their medians were closer to the medians of the reference rates (**Fig S5**). Modelling calibration also improved empirical estimates of a fungal growth rate for a non-parametric GP model. The CIs of the estimated growth rates included the CIs of the reference rates if calibration was modelled (GP-OD-calibration model), but not otherwise (GP-OD model) (**Fig S6**).

Our model treated the measured OD_600_ as distributed around a linear transform of the true fungal growth, while the Logistic-OD model without calibration inferred true fungal growth by fitting a logistic function directly to the OD_600_ data. Our inferred true fungal growth dynamics (**Fig 5b**) increased more rapidly compared to the measured OD_600_ data (**Fig 5c**). Hence, the hyphal growth rate estimated from our Logistic-OD-calibration model’s inferred true fungal growth dynamics was distinctly larger than hyphal growth rate estimated by the Logistic-OD model directly fit to the OD_600_ data.

These results suggest modelling calibration reduced the observed discrepancy between hyphal growth rates estimated from OD_600_ and directly measured data. Our Logistic-OD-calibration model, the Gompertz-OD-calibration model and the GP-OD-calibration model could all infer similar growth rates from all the data collection methods considered (OD_600_, HL, NC).

## Discussion

The need for surveillance of antifungal drug susceptibility is a pressing problem in medical mycology. However, investigating time- and morphology-specific antifungal modes of action using mechanistic modelling or estimating growth rates is currently hampered in clinical microbiology laboratories by arduous methodologies reliant upon sophisticated technologies. A potential solution is to use indirect measures of fungal growth, such as OD, for model fitting and fungal growth rate estimation. We found a logistic model could not be directly fit to OD_600_ data for low density fungal cultures and the estimated growth rate was distinct from reference rates estimated from directly measured data (**Fig 2**). To overcome these issues, we proposed a mathematical model that described the dynamic changes in fungal growth and incorporated calibration in the model. The model captured observed dynamics in OD_600_ growth data (**Fig 3**) and outperformed population growth models previously used in OD modelling in terms of RMSE and relative LPD when predicting held-out test data during CV (**Fig 4**). We then used the model to infer a hyphal growth rate from indirect OD_600_ data and a hyphal extension rate and growth rate from direct (HL, NC) measures (**Fig 5**). The fungal growth rate inferred from the OD_600_ data using our model was closer to those inferred from the direct measures of fungal growth compared to a reference logistic model that did not include calibration. Our model was able to infer growth rates with similar medians from all the collected data sets (OD_600_, HL, NC), indicating our estimates were robust to the data collection method.

Two similar attempts for modelling calibration have been previously made. A statistical model [43] was proposed to model a calibration process required for estimating unknown concentrations of any samples. Its success was demonstrated using cockroach allergens [43]. However, this model has not been used in conjunction with a population growth model to infer microbial growth rates to the best of our knowledge. Another study [38] proposed predicting microbial growth curves from OD by fitting population growth models to data that details the first time the OD reader can detect growth in wells initialised with different initial inocula. However, the method requires that the well’s maximum carrying capacity is known along with the microbial number at the first time the OD reader can detect growth or a pre-chosen OD value [38]. For filamentous fungi, we do not know the maximum carrying capacity of a well or the fungal concentration at the first OD detection time or any pre-chosen OD without the collection of calibration data.

In the wider microbiology community, calculating growth rates to characterise and compare growth dynamics of microbes, such as bacteria or yeast, in different conditions is common [38,40,41,44–52]. Growth rates have been estimated directly from OD using empirical methods with application to antimicrobial susceptibility testing [50,51] or using population growth models fit to OD data [48]. In addition, various methods to improve bacterial growth rate estimation from OD have been proposed, such as the use of GP models, where growth rates are calculated empirically post-inference [40,41,52]. However, none of these models [40,41,48,52] can be reused to estimate growth rates of filamentous fungi from OD because all the models either require OD data that has been pre-processed using collected calibration data or the models are directly fit to OD, with one study even highlighting OD calibration as a problem when estimating growth rates [44]. Calibration data that are available for microbes such as bacteria cannot be reused for filamentous fungi since OD values are sensitive to the size and shape of the microbe measured [34,35]. Moreover, we found that directly fitting models of true fungal growth to OD_600_ data resulted in model misfit and distinctly lower growth rates compared to growth rates estimated from direct measures for filamentous fungi.

For bacteria, non-parametric models have been preferentially used to estimate growth rates from OD because it has previously been recommended to calculate growth rates empirically instead of using population growth models [53]. In this study, we used non-parametric GP models (GP-OD and GP-OD-calibration) as reference models. The GP-OD-calibration model could successfully estimate an empirical growth rate that had CIs that included the CIs of reference rates estimated from direct measures of fungal growth, without the need for calibration data. However, we argue that our parametric Logistic-OD-calibration model is preferred for fungal growth modelling and growth rate estimation for the following three reasons. Firstly, we demonstrated that our model outperformed the non-parametric GP-based models both at fitting to and predicting fungal OD_600_ curves during CV (**Fig 4**). Secondly, our model benefits from explicit inclusion of a growth rate, as opposed to GP-based models that provide an empirical *post-hoc* estimation of the growth rate, because it increases biological interpretability. Lack of an explicit growth rate parameter in a model makes it more difficult to detect potential bias in the estimated growth rate arising from unknown external factors not accounted for in the model. Thirdly and finally, parametric models more readily allow for known fungal growth mechanisms to be included in the model than for non-parametric models, for example including different initial fungal inocula of an experiment in the model.

Previously, a non-mechanistic piecewise-linear model was fit to OD_600_ data to detect the number of change-points in the growth curves [54] that does not consider OD being an indirect measure of growth. We did not include it as a reference model in this study because it was not targeted at estimating growth rates or mechanistically modelling fungal growth. Fungal growth rates would need to be calculated empirically and subsequently compared between different experimental conditions. Different conditions may result in different numbers of estimated change-points, and difficult-to-collect calibration data would still be needed to estimate these growth rates.

In this paper, we assumed that the true latent growth follows a deterministic logistic growth model and that measured OD_600_ is a linear transform of true fungal growth (a linear calibration curve). Future research could investigate whether incorporating the natural stochasticity of fungal growth in our latent model would improve model fit. Our chosen logistic function for true fungal growth could also be expanded to investigate more complex morphology-specific growth mechanisms or to include antifungal activity. Methods have already been proposed to accelerate Bayesian inference of large-scale models whose linear transform is the only observed output [55], which would facilitate estimation of fungal growth rates should the expansion of our model dramatically increase computation time. Another interesting avenue for future research would be to explore more complex relationships between measured OD and fungal growth, since a linear calibration curve is not necessarily reflective of the true relationship between measured OD and fungal growth for large OD [34]. However, we anticipate that modelling a nonlinear calibration curve whilst retaining a parametric model would make key parameters of interest (e.g. fungal growth rates) difficult to infer because large differences in the parameter values may no longer correspond to measurable differences in observed OD.

In summary, this study proposed a mathematical model to infer a biologically interpretable and morphology-specific fungal growth rate from indirect OD_600_ data without the need for collecting calibration data and, hence, is accessible to all microbiology laboratories that collect OD_600_ data, rather than just specialist laboratories. The results shown here serve as the much-needed groundwork for investigating time- and morphology-specific changes to filamentous fungal growth using mathematical models with OD_600_ data. Future mechanistic models can use our presented model as a foundation to investigate mechanisms of antifungal action using only OD_600_ data by inferring and subsequently comparing growth rates of fungi in varying antifungal concentrations.

## Methods

### *In vitro* OD_600_ data of fungal growth

We obtained OD_600_ data from an experiment with wells containing different initial inocula of A1160P+[56] *A. fumigatus* (0, 2 × 10^2^, 2 × 10^3^, 2 × 10^4^ and 2 × 10^5^ spores) suspended in 200μl of fRPMI (0, 1, 10^1^, 10^2^ and 10^3^ [N/μl]) in three dimensional (3D) wells. The OD_600_ was recorded every hour for 24-25 hours for each well, for nine (three technical times three biological) replicates at wavelength 600nm by a BioTek OD reader. The wells with 0 spores were used to determine the background OD_600_ level.

### In vitro hyphal length data of fungal growth

We obtained *in vitro* HL data of fungal growth for A1160P+ [56] *A. fumigatus*. A 50μl sample from a spore suspension of 2 × 10^4^ spores/ml was added to 950μl of fRPMI in 3D wells to give an initial inoculum of 1 [N/μl] and images of the wells were captured every hour from 4 to 25 hours. The fRPMI growth media was chosen to match the growth media used in the OD_600_ experiment.

A hypha was chosen at random in the images for each replicate (*n* = 3) and the hyphal length was measured (μm) using ImageJ [36] until the single hypha could not be tracked anymore (18 hours) due to overcrowding or the hypha growing in a plane that is not in the field of vision of the image (e.g., upwards towards the camera).

### *In vitro* nuclear count data of fungal growth

A 100μl sample from a 10^5^ spores/ml suspension of fluorescent *A. fumigatus* spores (CEA10 strain (FGSC#1163) with the histone H1 tagged with GFP (PGpdA-H1-sGFP) [57]) was grown in 3D wells containing 900μl of fRPMI without phenol red (equivalent to an initial inoculum per well of 10^1^ [N/μl]) and images were taken every hour for 16 hours. A1160P+ and the histone tagged CE10 strain of *A. fumigatus* all derive from the CE10 strain [56,57]. fRPMI without phenol red was used in this experiment to avoid autofluorescence of the fRPMI interfering with counting fluorescent nuclei during data collection and we do not expect the exclusion of phenol red in the fRPMI media to alter *A. fumigatus* growth dynamics. Hyphae (*n* = 7) were followed and their number of nuclei per hypha were counted in a point-and-click fashion using the particle counting tool provided by ImageJ [36] until hyphae could no longer be followed (13 hours).

### Logistic-OD-calibration model

We describe the dynamics of true (latent) fungal concentration, *f*(*t*) [N/μl], at time *t* by a logistic function that begins growth at *t* = *τ* hours,

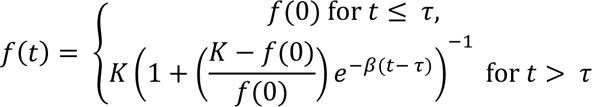

where *β* is the hyphal growth rate, *K* is the carrying capacity and *τ* represents the time delay for the fungal growth to begin after inoculation. The initial condition, *f*(0), is the corresponding initial fungal inoculum used in the OD_600_ experiment.

We model the measured OD_600_, *y*_*t*_, at time *t*, as distributed around a linear transform of the latent fungal growth with multiplicative noise that is proportional to the OD_600_ value (as we expect the noise to increase for high OD_600_ values without exceeding measured OD_600_ for small OD_600_ values),

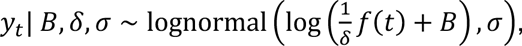

where δ (>1) is a proportionality constant that represents the proportion of light absorbance, *σ* is the scale of the measurement noise and *B* is the mean basal OD_600_ reading corresponding to the mean OD_600_ of the background fRPMI media (background correction). Priors were chosen for each of the parameters (*B*, *δ*, *σ*, *β*, *K*, *τ*) (details in Supplementary) and a prior sensitivity analysis was conducted for the growth rate *β* (**Fig S7**), where Bayesian inference for the Logistic-HL model with the less informative prior was conducted with adapt_delta set to 0.99 to ensure there were no signs of non-convergence. The model is reduced to a logistic function with multiplicative noise, *y*_.,*t*_∼ *lognormal*(*log*(*f*(*t*)), *σ*.), when we fit the model to direct measurements of fungal growth (HL data, *y*_*H*,*t*_, and the NC data, *y*_*N*,*t*_), where a different noise scale (*σ*.) is inferred for the HL and the NC data, *σ*_*H*_ and *σ*_*N*_, respectively.

### Models considered

We considered the following fifteen reference models in addition to our Logistic-OD-calibration model (Table 1).

**Table 1:**
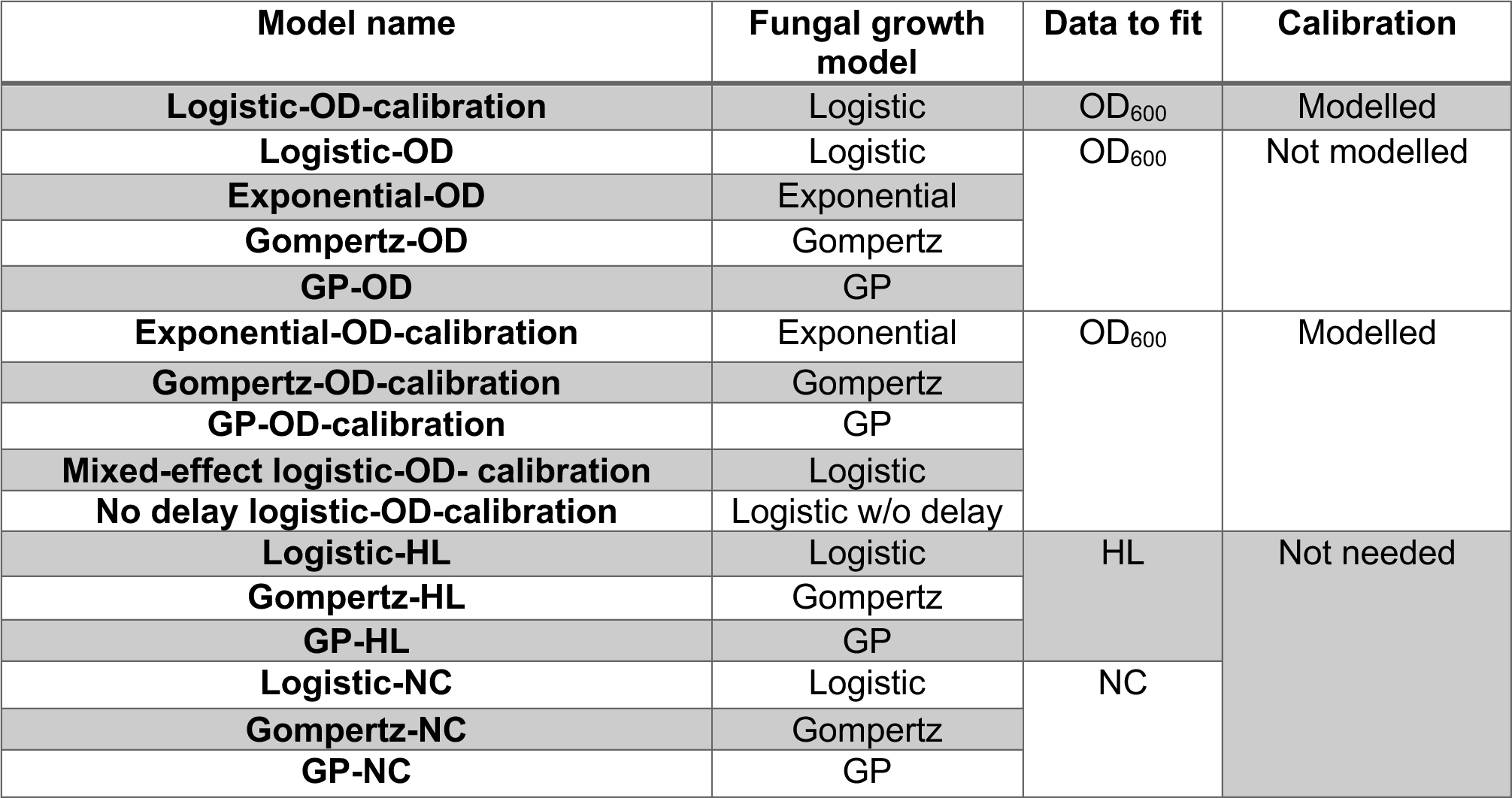
Our model and the reference models considered in this study.

The models have three defining aspects: the fungal growth model, *f*(*t*), (logistic, Gompertz, GP, exponential or logistic with no delay), the data they are fit to (OD_600_, HL or NC) and whether calibration is included in the model (modelled, not modelled or not needed). We considered the following four underlying fungal growth models in conjunction with three ways to model the measured OD_600_, depending on whether calibration is modelled, not modelled, or not needed.

#### 1. Fungal growth models

**Logistic** 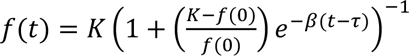 for *t>τ*, with parameters *K*, *β* and *f*(0) being the carrying capacity, the hyphal growth rate and a parameter for the initial OD_600_ value or the known initial fungal inoculum when calibration is included in the model. For the logistic with no delay, *τ* = 0.

**Gompertz** 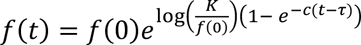, with *K*, *β*, and *f*(0) being the carrying capacity, the initial growth rate and a parameter for the initial OD_600_, respectively. When calibration is included in the model the known initial fungal inoculums of the OD_600_ experiment are used instead of a parameter for the initial OD_600_ value.

**GP** *f*(*t*), where *f*(*t*) is a GP model [58] for the log-transformed OD_600_ data with a zero mean GP prior and an exponentiated quadratic kernel, *K*(***t***, ***t***), which is an *n* by *n* matrix whose (*i*, *j*)-th element is 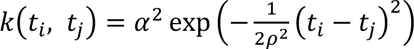 [58]. We assumed Gaussian noise on the log OD_600_ centred around the GP, *f*(*t*), that had a scale of *σ*, log *y*_*t*_ ∼ *N*(*f*(*t*), *σ*).

As described in Swain *et al.* [40], the growth rate was calculated by taking the maximum value of the time derivative of the fitted GP, which is analogous to the maximum time derivative of the logarithm of the growth curve. We did not apply a media correction to the data before fitting to ensure strictly positive data for calculating logarithms. This does not alter the inferred growth rate as derivatives are invariant to translations.

**Exponential** *f*(*t*) = *f*(0)*e*^*β*(*t*− *τ*)^ for *t> τ*, where *β* is the growth rate and *f*(0) specifies the parameter for the initial fungal inoculum, which is known when calibration is included in the model.

#### 2. Modelling calibration

When calibration is modelled, OD_600_ is assumed to be distributed around a linear transform of the (latent) true fungal growth model with multiplicative noise that has a scale of *σ*, *y*_*t*_| *B*, *δ*, *σ* ∼ 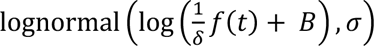, where *B* represents the OD_600_ of the background fRPMI media and 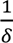 is the proportionality constant. To avoid negative values for the true latent fungal growth when using the GP fungal growth model (GP-OD-calibration model), we model *f*(*t*) as a GP for the log-transformed latent fungal growth, 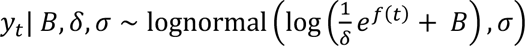. The calculated growth rate is again the maximum value of the time derivative of the fitted GP, which is analogous to taking the maximum value of the time derivative of the logarithm of the latent growth curve. Hence, the growth rate estimated from the GP-OD-calibration model is comparable to growth rate estimated using GP-OD where calibration is not modelled.

In the mixed-effect logistic-OD-calibration model, we assume random effects on the background media parameter *B*. We model a background media OD_600_ value, *B*^j^, for the *j*-th replicate 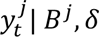, 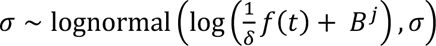, and assume that the *B*^j^ are distributed around the sample mean, 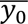, of the observed blanks (wells that had an inoculum size of 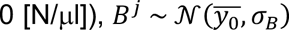. The sample mean is calculated by 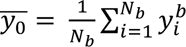, where 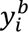 denotes the measured OD_600_ values of the blanks and *N*_*b*_ is the total number of data points of the blanks.

If the calibration is not modelled, either OD_600_ is assumed to be distributed around the fungal growth model with multiplicative noise with a scale of *σ*, *y*_*t*_ ∼ lognormal(log(*f*(*t*)), *σ*), or the fungal growth model will specify its own model for the OD_600_ data (as in the GP fungal growth model).

Including calibration in the model is not needed if a model is fit to the directly measured (HL and NC) data.

### Model inference

For all models considered, we sampled posterior distributions using the Hamiltonian Monte Carlo (HMC)-based No-U-Turn sampler (NUTs) [59] provided by RStan [60]. The models were run for 2000 iterations using 50% for warm-up for four chains. The Logistic-OD model was run with the control parameter adapt_delta set to 0.99 during model fitting to ensure there were no signs of non-convergence. The GP-OD-calibration model was run for 4000 iterations with adapt_delta set to 0.99 during model fitting to ensure there were no signs of non-convergence. The Logistic-OD-calibration model was also run for 4000 iterations with adapt_delta at 0.99 for sampling only during prior predictive checks only to ensure no signs of non-convergence. The convergence was monitored using the Gelman-Rubin metric 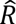 [61], where 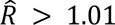 was used as a heuristic to diagnose a lack of convergence. All models were developed and checked using a full Bayesian Workflow through prior, fake data and posterior predictive checks [62]. Prior predictive and fake data checks were conducted to confirm that priors reflected expert knowledge about fungal OD_600_ values (**Fig S8**) and that all the parameters could be estimated using the model with these priors from fake data (**Fig S9**). Posterior predictive checks were conducted to assess the models’ potential ability to explain the observed data. We fit each of the models to the OD_600_ fungal growth data (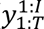 where *I* is the total number of replicates and *T* is the final time the OD_600_ is measured (=25 hours)) and sampled replicates, 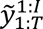, from its posterior predictive distributions, 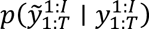. We then visually checked if 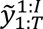 demonstrate and a sufficiently close resemblance to 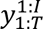.

### Predictive performance

We assessed the performance of the models on predicting entire held-out time-series of replicates using 5-fold CV. The training-testing data split was stratified by the replicates. Model fit or predictive performance was assessed using the following two metrics:

1. Relative log (pointwise) predictive density (LPD): 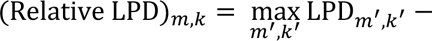 − LPD_*m*,*k*_ for the *m*-th model and the *k*-th CV fold, where the LPD_*m*,*k*_ is the LPD averaged over the testing replicates in fold *k*, *k*_test_. The LPD is 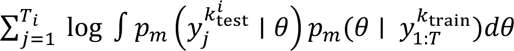 [63] for the *i*-th replicate in *k*_test_ measured until a time *T_i_*, where *k*_train_ and *k*_test_ are the training and testing replicate indices, respectively, within the *k*-th fold. The LPD is calculated by 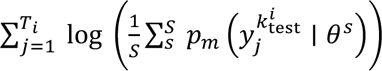 [63], where *s* is the index of a sample from the posterior, 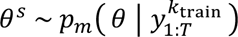, *S* is the total number of posterior samples and *j* is a given data point of the *i*-th replicate in *k*_test_.
2. The root mean squared error (RMSE): RMSE_*m*,*k*_ is calculated between the model’s predictions, 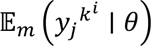, and the observed data, 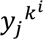, at the *j*-th data point in the *i*-th replicate for the *k*-th fold, 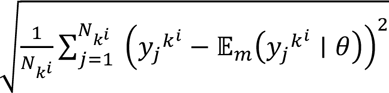, and averaged over the replicates, where *N*_*k*_^*i*^ is the total number of data points for replicate *i* in the test or training data set in the CV fold *k*.

The RMSE of an intercept model (mean of all OD_600_ values) was included during model comparison as a baseline (**Fig 4**).

## Contributions

**T.H.** – Conceptualization, Investigation, Data Curation, Formal Analysis, Funding Acquisition, Methodology, Software, Validation, Visualization, Writing – Original Draft Preparation. **N.M.** – Data Curation, Investigation. **E.B.** – Conceptualization, Funding Acquisition, Resources, Writing – Review & Editing. **R.J.T.** – Conceptualization, Project Administration, Funding Acquisition, Supervision, Writing – Review & Editing.

## Data availability statement

All code used for model fitting and plotting is available on a GitHub repository at https://github.com/tah17/Inferring-fungal-growth-rates-from-OD-data.

## Financial Disclosure Sentence

This research was funded in whole, or in part, by the Wellcome Trust [Grant number 215358/Z/19/Z to TH], https://wellcome.org/. For the purpose of open access, the author has applied a CC BY public copyright licence to any Author Accepted Manuscript version arising from this submission. This project was partly funded by National Centre for the Replacement, Refinement and Reduction of Animals in Research (NC3Rs) Studentship (NC/P00217X/1 to N.M. and R.J.T.), https://nc3rs.org.uk/. E.B acknowledges support from the MRC Centre for Medical Mycology at the University of Exeter (MR/N006364/2 and MR/V033417/1). Work in the laboratory of E.B. was funded by MRC project grants MR/M02010X/1, MR/S001824/1, and MR/L000822/1 and by a Biotechnology and Biological Sciences Research Council (BBSRC) project grant BB/V017004/1. The funders had no role in study design, data collection and analysis, decision to publish, or preparation of the manuscript.

## Acknowledgements

This research was funded in whole, or in part, by the Wellcome Trust [Grant number 215358/Z/19/Z]. For the purpose of open access, the author has applied a CC BY public copyright licence to any Author Accepted Manuscript version arising from this submission.

