## Supplementary Information for "Inferring fungal growth rates from optical density data"

Tara Hameed[1], Natasha Motsi[1], Elaine Bignell[2] and Reiko J. Tanaka[1]
1. Department of Bioengineering, Imperial College London, London, United Kingdom. 2. Medical Research Council Centre for Medical Mycology at The University of Exeter, University of Exeter, Exeter, United Kingdom.

**Supplementary Information**

### A Supplementary figures and tables

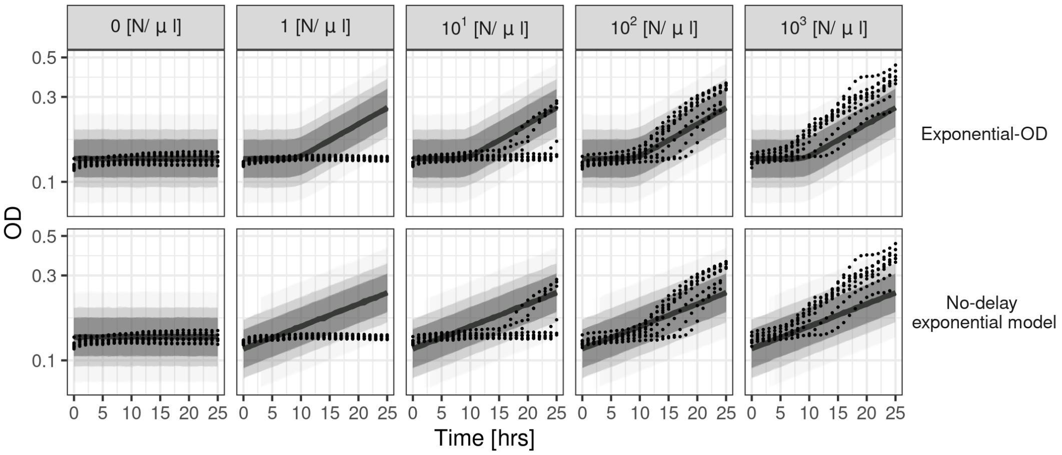


**Fig. S1**: **Exponential models fit to $OD_{600}$ data**. Posterior predictive distribution of the Exponential-OD model (top) and an exponential model with no delay (bottom) fit to the $OD_{600}$ data (black dots) with all the initial fungal inocula (0, 1, $10^1$, $10^2$, $10^3$ [N/$\mu$l], left to right). Solid lines and shades are medians and 95%, 80%, 60% credible intervals, respectively. The y-axis is in $\log_{10}$ scale.

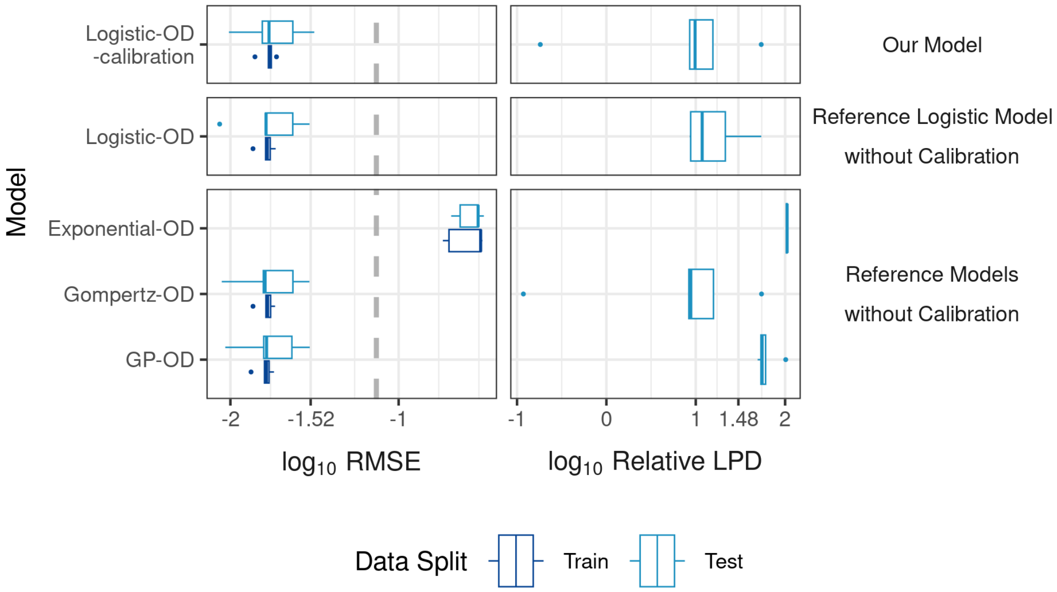


**Fig. S2**: **Model fit and prediction using only high fungal inoculum ($10^3$ [N/$\mu$l]) $OD_{600}$ data**. Model fit and prediction are quantified by RMSE (3.s.f.) averaged over training (dark blue) and testing (light blue) $OD_{600}$ replicates, respectively, for each fold in 5-fold cross validation stratified by replicates (left). The dotted grey line indicates the RMSE of an intercept model. Model prediction is quantified by Relative LPD ((maximum LPD) - LPD) (3.s.f.) with LPD averaged over test replicates (right). Lower values indicate better fitting and predictive performance for both metrics.

**Table S1**: **The relative LPD and RMSE during CV for GP-OD, GP-OD-calibration, Logistic-OD-calibration and No-delay logistic-OD-calibration models.** Median, 25% and 75% quantiles of relative LPD and RMSE averaged over the replicates in each fold for the GP-OD, GP-OD-calibration, Logistic-OD-calibration and No-delay logistic-OD-calibration models during cross validation. Differences in the values between the models (GP vs GP-OD-calibration and Logistic-OD-calibration vs No-delay logistic-OD-calibration) are highlighted in bold.

| **Metric** | **Data split** | **Model** | **Median (5.s.f.)** | **25% Quantile (4.s.f.)** | **75% Quantile (4.s.f.)** |
|---|---|---|---|---|---|
| Relative LPD | Testing | GP | 89.4**56** | 88.**91** | 90.**25** |
| | | GP-OD-calibration | 89.4**58** | 88.**87** | 90.**21** |
| | | Logistic-OD-calibration | 23.**760** | 22.1**8** | 25.6**7** |
| | | No-delay logistic-OD-calibration | 23.**711** | 22.1**7** | 25.6**6** |
| RMSE | Testing | GP | 0.042**728** | 0.04**097** | 0.04**304** |
| | | GP-OD-calibration | 0.042**666** | 0.04**109** | 0.04**318** |
| | | Logistic-OD-calibration | 0.0186**82** | 0.0186**3** | 0.0225**5** |
| | | No-delay logistic-OD-calibration | 0.0186**68** | 0.0186**1** | 0.0225**3** |
| | Training | GP | 0.040**252** | 0.0399**5** | 0.0406**0** |
| | | GP-OD-calibration | 0.040**363** | 0.0399**8** | 0.0406**2** |
| | | Logistic-OD-calibration | 0.0186**80** | 0.018**40** | 0.018**69** |
| | | No-delay logistic-OD-calibration | 0.0186**44** | 0.018**37** | 0.018**66** |

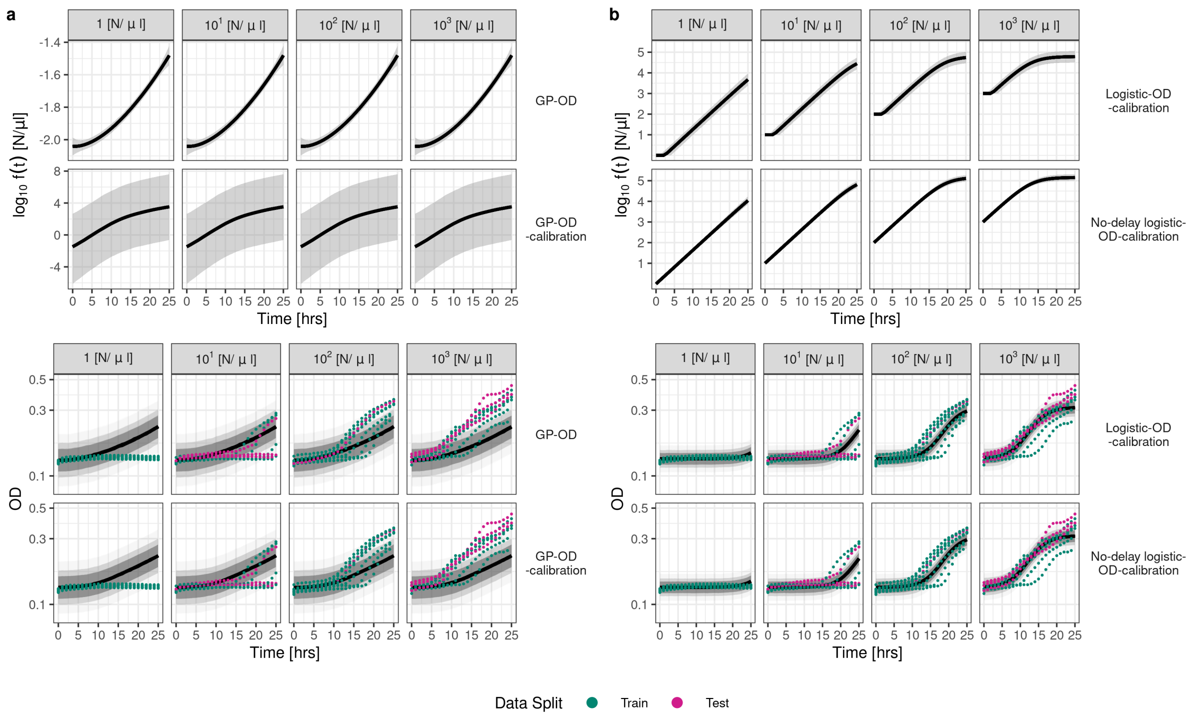


**Fig. S3**: **GP-based and "Logistic-OD-calibration"-based model fits in cross validation fold 3**. *Top*: The fungal growth, $f(t)$ [N/$\mu$l] ($\log_{10}$ scale), inferred from models fit to training data in cross validation fold 3 ($OD_{600}$ of wells inoculated with (1, $10^1$, $10^2$, $10^3$ [N/$\mu$l], left to right). Solid lines and shades are medians and 95% credible intervals, respectively. *Bottom*: Model fit (posterior predictive distribution, medians shown in solid black line and 95%, 80%, 60% credible intervals shaded, y-axis in $\log_{10}$ scale) to training $OD_{600}$ data in fold 3 (*green*) visually compared to the testing data (*pink*) for the GP-OD and GP-OD-calibration models (**a**) and Logistic-OD-calibration and No-delay logistic-OD-calibration models (**b**).

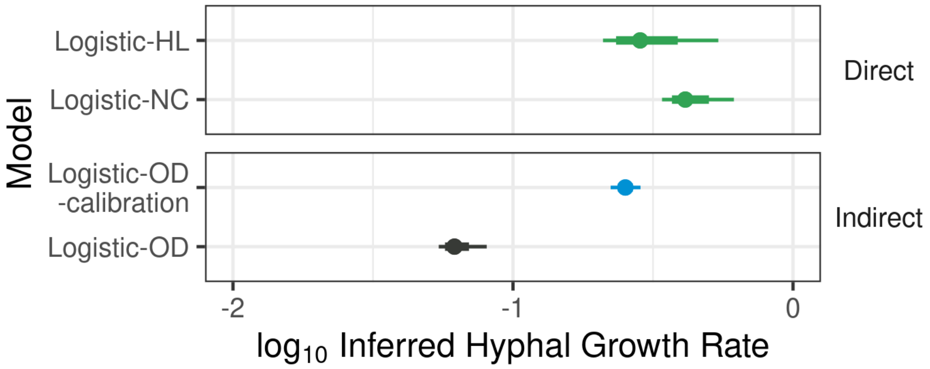


**Fig. S4**: **Hyphal growth rates inferred using only high fungal inoculum ($10^3$ [N/$\mu$l]) $OD_{600}$ data**. Medians (dot), 95% (thin line) and 80% (thick line) credible intervals of the reference hyphal growth rates estimated from logistic models fit to direct data (Logistic-HL and Logistic-NC fit to hyphal length and nuclear count data, respectively) and indirect $OD_{600}$ data of the high initial fungal inoculum ($10^3$ [N/$\mu$l]). The growth rate inferred from $OD_{600}$ data using our Logistic-OD-calibration model (80% and 95% CIs [0.232, 0.273] and [0.223, 0.285], respectively) (blue) is closer to the reference growth rates (green) than the rate from the Logistic-OD model (black). All the inferred rates are shown on a $\log_{10}$ scale.

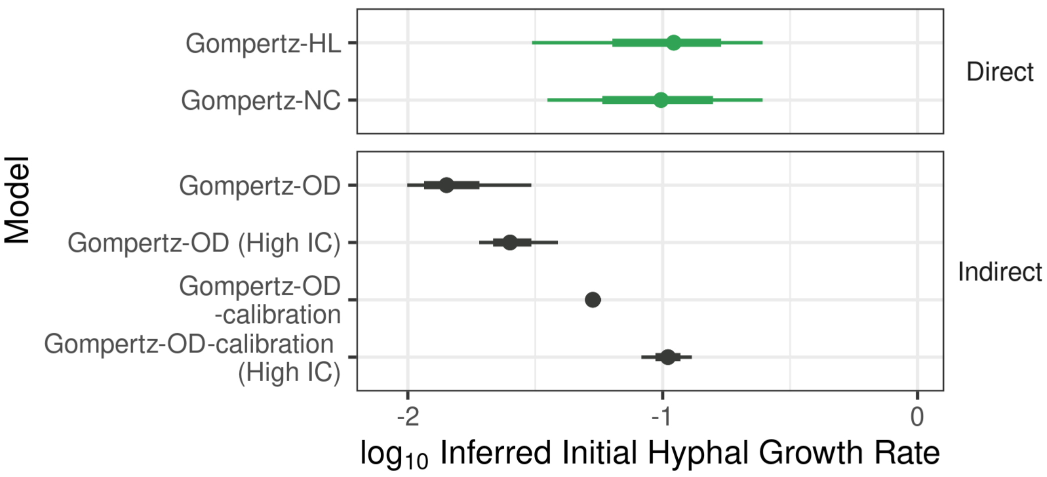


**Fig. S5**: **Fungal growth rates inferred by Gompertz models.** Initial fungal growth rates estimated by Gompertz models fit to directly measured data sets, hyphal length and nuclear count data, (reference rates, green) and indirect $OD_{600}$ data (black) of all the fungal inocula or just using the inoculum of $10^3$ [N/$\mu$l] (High initial condition (IC)). The Gompertz-OD model can only infer a growth rate with overlapping 95% credible intervals to the reference rates when fit to only the high initial inoculum data. However, the Gompertz-OD-calibration model could infer growth rates with credible intervals that were included in both reference rates' credible intervals using both all (Gompertz-OD-calibration model, 95% credible interval of the $\log_{10}$ growth rate is [-1.30, -1.24] to 3.s.f) or just high initial inoculum (Gompertz-OD-calibration model (High IC)) $OD_{600}$ data. All growth rates are shown on a $\log_{10}$ scale.

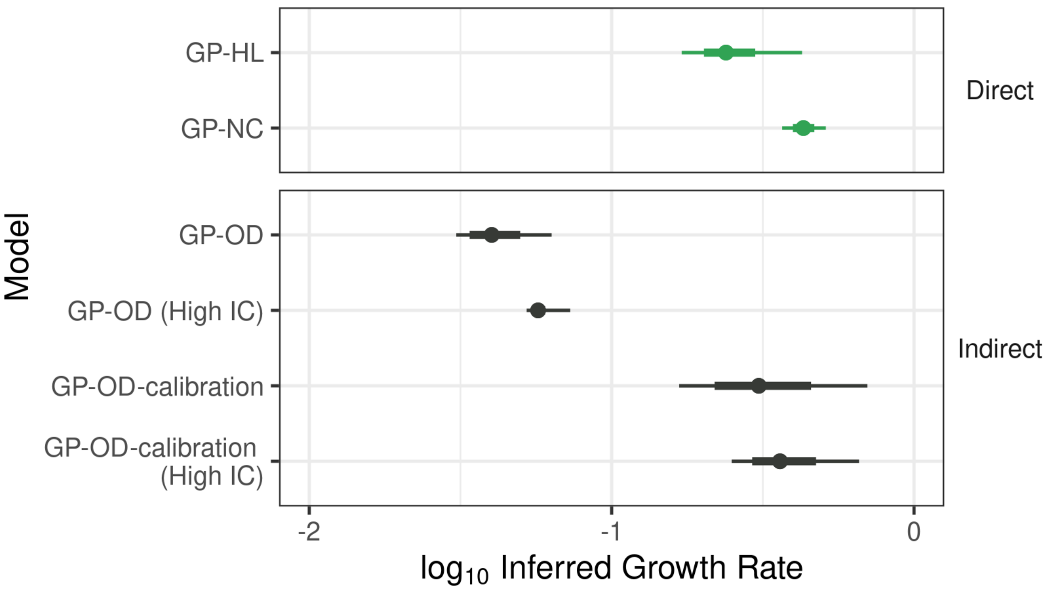


**Fig. S6**: **Fungal growth rates inferred by GP models.** Fungal growth rates estimated by GP-based models fit to directly measured data sets, hyphal length and nuclear count data, (reference rates, green) and indirect $OD_{600}$ data (black) of all the fungal inocula or just using the inoculum of $10^3$ [N/$\mu$l] (High initial condition (IC)). The GP-OD model that does not include calibration fails to infer similar growth rates to the reference growth rates derived from the GP-HL and GP-NC models when using all or just high initial inoculum $OD_{600}$ data. However, the GP-OD-calibration model that includes calibration could infer growth rates with credible intervals that overlapped with the reference rates' credible intervals using all (GP-OD-calibration model) or just high initial inoculum (GP-OD-calibration model (High IC)) $OD_{600}$ data. All growth rates are shown on a $\log_{10}$ scale.

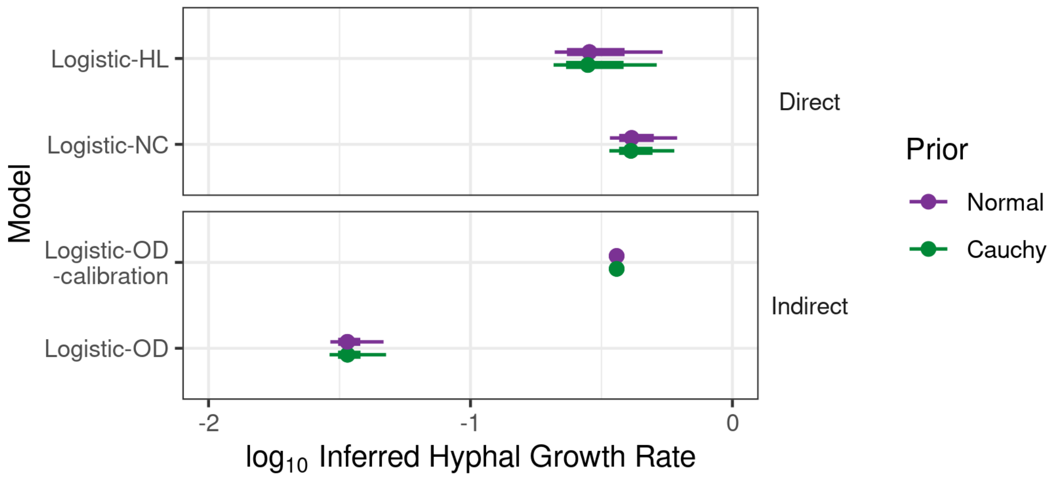


**Fig. S7**: **Sensitivity of inferred hyphal growth rates to prior choice.** Medians (dot), 95% (thin line) and 80% (thick line) credible intervals of the hyphal growth rates from logistic models with priors for the growth rates of the original half Normal priors (purple) used to generate the results vs less informative half Cauchy priors (green). Growth rates were inferred from logistic models fit to the direct data (Logistic-HL and Logistic-NC fit to hyphal length and nuclear count data, respectively) and indirect $OD_{600}$ data (Logistic-OD, Logistic-OD-calibration). The estimated growth rates' median and credible intervals do not depend on the prior choice apart from the Logistic-HL's 95% credible intervals. Even so, all growth rates inferred using our Logistic-OD-calibration model have credible intervals that are included in the credible intervals of the reference growth rates. All the inferred rates are on a $\log_{10}$ scale.

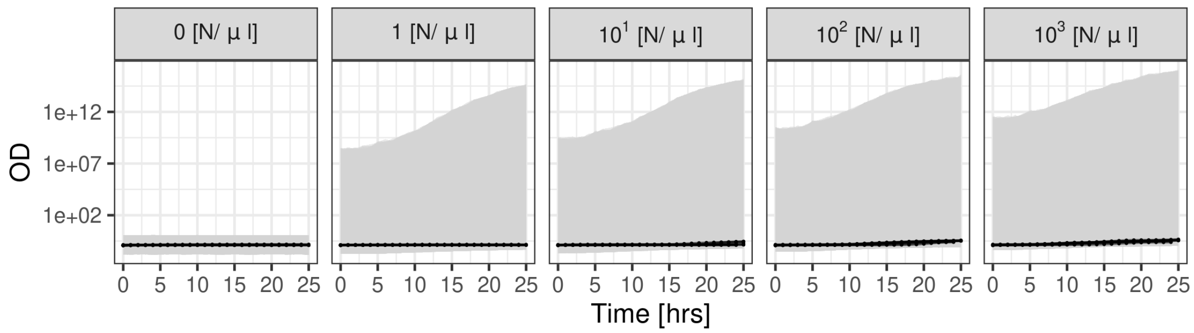


**Fig. S8**: **Prior predictive check of Logistic-OD-calibration model.** 95% credible interval (grey) of the prior predictive distribution for our Logistic-OD-calibration model compared to the observed $OD_{600}$ data (black dots, replicates joined by lines). The prior predictive distribution's density covers the $OD_{600}$ data, only includes positive $OD_{600}$ values and allows for sufficiently large $OD_{600}$ values. The y-axis is in $\log_{10}$ scale.

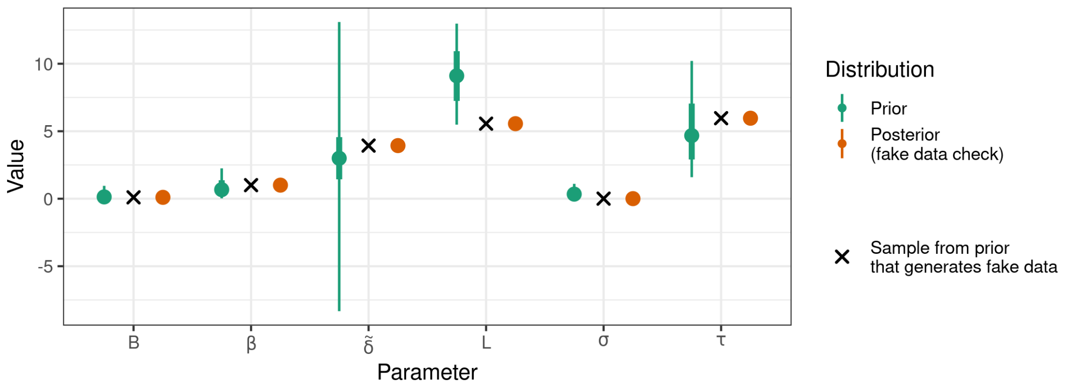


**Fig. S9**: **Results of fake data check for our Logistic-OD-calibration model**. Medians (dots) and 95% and 80% credible intervals (thin and thick lines, respectively) of priors (green) and estimated parameters obtained by fitting our model to a fake data set (orange) generated by sampling arbitrary parameter values from our Logistic-OD-calibration model's prior (black crosses). The parameters' 95% credible intervals estimated (orange) ([$1.0188 \times 10^{-1}$, $1.0199 \times 10^{-1}$], [1.0057, 1.0064], [3.9346, 3.9371], [5.5588, 5.5614], [$5.7817 \times 10^{-3}$, $6.2751 \times 10^{-3}$], [5.9606, 5.9677], 5.s.f.) include the initially chosen parameter values ($1.0198 \times 10^{-1}$, 1.0063, 3.9360, 5.5605, $6.2294 \times 10^{-3}$, 5.9659) to 5.s.f. (crosses), as desired.

### B Model priors

#### B.1 Logistic-OD-calibration model

Optical density measured at 600nm ($OD_{600}$) data, $y_t$, at a time, $t$, is modelled as a linear transform of true fungal growth, $f(t)$ [N/$\mu$l], with proportionality constant, $\frac{1}{\delta}$, and an offset parameter, $B$, representing the average basal $OD_{600}$ level. $f(t)$ is a logistic growth model that begins growth after a time $t > \tau$ [hours (h)] (referred to as a "delay" hereafter) with a (hyphal) growth rate $\beta$:

$$y_t|B, \delta, \sigma, \beta, K, \tau \sim \text{lognormal}\left(\log\left(B + \frac{f(t)}{\delta}\right), \sigma\right)$$

$$\text{where } f(t) = \begin{cases} f(0) \text{ for } t \leq \tau, \\ K\left(1 + \left(\frac{K - f(0)}{f(0)}\right) e^{-\beta(t-\tau)}\right)^{-1} \text{ for } t > \tau. \end{cases}$$

where $\sigma$ is the scale of the lognormal distribution. The priors of the parameters in the model are:

- $\tau$ [h]: time before fungal germination from spores to hyphae, which is positive and was observed to be around 5 [h] in the hyphal length (HL) and nuclear count (NC) data; $\tau \sim \text{Gamma}(5, 1)$.
- $\beta$ [$h^{-1}$]: hyphal growth rate. We know this to be positive and previous literature has found the proliferation rate to be around 0.3 [$h^{-1}$] [1]. Hence, we use $\beta \sim \mathcal{N}^+(0, 1)$, where $\mathcal{N}^+(0, 1)$ denotes a standard normal distribution truncated to the domain $[0, \infty)$.
- $K$: average well carrying capacity (maximum capacity). A prior is placed on a transform, $L = \log_{10}(K)$, $L \sim \mathcal{N}(9, 2)$, where the normal distribution is truncated to the domain $[\log_{10}(10^3), \infty)$. The highest initial inoculum ($f(0)$) used is $10^3$ [N/$\mu$l] and we expect the wells' carrying capacity to cover values sufficiently higher than, but never be lower than, this.
- $B$: mean $OD_{600}$ of the background fungal (f) Roswell Park Memorial Institute (RPMI) media. We assume that the true mean of the background fRPMI will be distributed around the sample mean of $OD_{600}$ values of the fRPMI only wells (blanks): $B \sim \text{lognormal}(\log(\bar{y}_0), 1)$, where $\bar{y}_0 = \frac{1}{N_b}\sum_{i=1}^{N_b} y_i^b$ and $y_i^b$ denotes the $OD_{600}$ values of the blanks and $N_b$ is the total number of data points of the blanks.
- $\sigma$: multiplicative noise scale. We do not expect the noise to exceed 100% of the observed value on average; $\sigma \sim \mathcal{N}(0, 0.5)$.

- $\delta$: proportionality constant. A prior is placed on a transform of $\tilde{\delta} = \log_{10}(\delta)$, $\tilde{\delta} \sim \mathcal{C}\left(\log_{10}(\max(f(0))), 1\right)$, where $\mathcal{C}$ is the Cauchy distribution. The highest initial inoculum size of $10^3$ [N/$\mu$l] results in $OD_{600}$ values that are indistinguishable from $OD_{600}$ of the blanks, so we place higher prior mass on values for the proportionality constant that are higher than the maximum initial inoculum.

The dependence of the hyphal growth rate on the prior choice is checked by repeating the inference but using the less informative prior of $\beta \sim \mathcal{C}^+(0,1)$ (**Fig. S7**) and the prior distributions for all the parameters are checked to be reasonable using a prior predictive check (**Fig. S8**).

When $f(0) = 0$ the following model is used

$$y_{t,b}|B,\sigma \sim \text{lognormal}\left(\log(B),\sigma\right)$$

where $y_{t,b}$ is the $OD_{600}$ data of the blanks.

#### B.2 Reference models

##### B.2.1 Logistic-OD

$OD_{600}$ data, $y_t$, at a time, $t$, is modelled as centred around logistic fungal growth, $f(t)$ [N/$\mu$l], that begins growth after a time $t > \tau$ [h]. The logistic function has a growth rate $\beta$ and carrying capacity $K$.

$$y_t|\sigma,\beta,K,\tau,f(0) \sim \text{lognormal}\left(\log\left(f(t)\right),\sigma\right)$$
$$\text{where } f(t) = \begin{cases} f(0) \text{ for } t \le \tau, \\ K\left(1 + \left(\frac{K - f(0)}{f(0)}\right)e^{-\beta(t-\tau)}\right)^{-1} \text{ for } t > \tau. \end{cases}$$

The following parameters have the same priors as in the Logistic-OD-calibration model (Section B.1); $\tau$, $\beta$, $\sigma$ and $B$. The parameters that have different priors are listed below with their corresponding prior distributions:

- $K$: the carrying capacity. This logistic model is fit directly to the $OD_{600}$ data, so we assume that the average carrying capacity is centred around 1 (a large value for $OD_{600}$ data), $K \sim \mathcal{N}^+(1,1)$;
- $f(0)$: the initial $OD_{600}$ value. We place a $f(0) \sim \text{lognormal}(\log(1), 0.5)$ prior on this parameter.

As in the above section (Section B.1), the dependence of the fungal growth rate on the prior choice is checked by repeating the inference but using the less informative prior of $\beta \sim \mathcal{C}^+(0,1)$ (**Fig. S7**). Finally, the $OD_{600}$ data of the blanks, $y_{t,b}$, ($f(0) = 0$) is modelled using $y_{t,b}|B,\sigma \sim \text{lognormal}\left(\log(B),\sigma\right)$, as in Section B.1.

##### B.2.2 Exponential-OD

$OD_{600}$ data, $y_t$, at a time, $t$, is modelled as centred around an exponential function, $f(t)$ [N/$\mu$l], that begins growth after a time $t > \tau$ [h], where $\beta$ and $f(0)$ are the hyphal growth rate and a parameter for the initial $OD_{600}$, respectively.

$$y_t|\sigma,\beta,\tau,f(0) \sim \text{lognormal}\left(\log f(t),\sigma\right)$$
$$\text{where } f(t) = \begin{cases} f(0) \text{ for } t \le \tau, \\ f(0)\exp\left\{\beta(t-\tau)\right\} \text{ for } t > \tau. \end{cases}$$

The growth rate, $\beta$, the delay, $\tau$, the parameter for the initial $OD_{600}$, $f(0)$, and the scale of the noise, $\sigma$, are given the same priors as the priors of the growth rate, $\beta$, delay $\tau$, the initial $OD_{600}$, $f(0)$, and the noise scale, $\sigma$, in the Logistic-OD model (Section B.2.1), respectively. The $OD_{600}$ data of the blanks, $y_{t,b}$, is again modelled using $y_{t,b}|B,\sigma \sim \text{lognormal}\left(\log(B),\sigma\right)$, as in Section B.1, and the prior for $B$ is kept the same as in Section B.1.

##### B.2.3 Gompertz-OD

$OD_{600}$ data, $y_t$, at a time, $t$, is modelled as centred around Gompertz function that begins growth after a time $t > \tau$ [h], $f(t)$ [N/$\mu$l], where $K$, $c$, and $f(0)$ being the carrying capacity, the initial growth rate and a parameter for the initial $OD_{600}$.

$$y_t|\sigma, c, K, \tau, f(0) \sim \text{lognormal}\left(\log f(t), \sigma\right)$$
$$\text{where } f(t) = \begin{cases} f(0) \text{ for } t \leq \tau, \\ f(0) \exp\left\{\log\left(\frac{K}{f(0)}\right)\left(1 - e^{-c(t-\tau)}\right)\right\} \text{ for } t > \tau. \end{cases}$$

The carrying capacity, $K$, the initial growth rate, $c$, the delay, $\tau$, the parameter for the initial $OD_{600}$, $f(0)$, and the noise scale, $\sigma$, are given the same priors as the priors of the carrying capacity, $K$, growth rate, $\beta$, the delay, $\tau$, the initial $OD_{600}$, $f(0)$, and the noise scale $\sigma$, in the Logistic-OD model (Section B.2.1), respectively. Finally, the $OD_{600}$ data of the blanks, $y_{t,b}$, is modelled using $y_{t,b}|B, \sigma \sim \text{lognormal}\left(\log(B), \sigma\right)$, with the prior for $B$ kept the same as in Section B.1.

### B.2.4 GP-OD

We model the log-transformed $OD_{600}$ data using a Gaussian Process (GP) [2]: $f(t)$, where $\sigma$ is the scale of the observed noise of the log $OD_{600}$.

$$\{\log y_t\}|\alpha, \rho, \sigma \sim \mathcal{N}\left(f(t), \sigma\right)$$

The GP is equipped with a zero mean GP prior and the exponentiated quadratic kernel, $K(\boldsymbol{t}, \boldsymbol{t})$, where the $(i, j)$-th element is defined by

$$\text{k}\left(t_i, t_j\right) = \alpha^2 \exp\left(-\frac{1}{2\rho^2}\left(t_i - t_j\right)^2\right)$$

The following priors are used for the parameters in the GP-OD model:

- $\rho$, the length scale of the GP. We expect $OD_{600}$ values to change as quickly as 10 hours post inoculation and should not have a length scale that exceeds the observed maximum time[1] ($t = 25$ hours): $\rho \sim \text{Gamma}(10, 1)$.
- $\alpha$, the output scale of the GP. The $OD_{600}$ data generally falls in the range of $[0, 1]$, hence we do not expect the log $OD_{600}$ to widely vary $\alpha \sim \mathcal{N}(0, 0.5)$;
- $\sigma$: the noise scale of the log data. We do not expect the noise to exceed 100% of the $OD_{600}$ values on average; $\sigma \sim \mathcal{N}(0, 0.5)$.

For the data from the blanks, a separate GP is fit to the log of the blank data, $\{\log y_{t,b}\}|\alpha_b, \rho_b, \sigma \sim \mathcal{N}\left(f_b(t), \sigma\right)$. The priors for $\alpha_b$ and $\rho_b$ are the same as the ones given for $\alpha$ and $\rho$ above.

In the GP-OD model, it is possible to directly evaluate the analytical posterior of the derivative, $\boldsymbol{g}$, of the GP, $\boldsymbol{f}$, evaluated at time points, $\boldsymbol{t}^d$, where we want to predict the derivative, following Swain *et al.* [3]:

$$\boldsymbol{g}|\boldsymbol{t}, \boldsymbol{y}, \boldsymbol{t}^d \sim \text{MVN}(\mathbb{E}(\boldsymbol{g}), \text{cov}(\boldsymbol{g}))$$
$$\mathbb{E}(\boldsymbol{g}) = \partial_1 K(\boldsymbol{t}^d, \boldsymbol{t})[K(\boldsymbol{t}, \boldsymbol{t}) + \sigma^2 I_n]^{-1}\boldsymbol{y}$$
$$\text{cov}(\boldsymbol{g}) = \partial_1\partial_2 K(\boldsymbol{t}^d, \boldsymbol{t}^d) - \partial_1 K(\boldsymbol{t}^d, \boldsymbol{t})[K(\boldsymbol{t}, \boldsymbol{t}) + \sigma^2 I_n]^{-1}[\partial_1 K(\boldsymbol{t}^d, \boldsymbol{t})]^{\text{T}}$$

where MVN is a multivariate normal distribution and $n$ is the number of observed time points, $\boldsymbol{t}$. We take the growth rate as the maximum value of the derivative, $\boldsymbol{g}$, evaluated at time points, $\boldsymbol{t}$.

[1] As advised in Betancourt, Michael (2020). Robust Gaussian Process Modeling, 2020. Retrieved from https://github.com/betanalpha/knitr_case_studies/tree/master/gaussian_processes, commit e10083abbcdb65c745f840ab9d2da58229fa9af3

##### B.2.5 Exponential-OD-calibration

As in Section B.1, $OD_{600}$ data, $y_t$, at a time, $t$, is modelled as a linear transform of latent fungal growth, $f(t)$ [N/$\mu$l], with proportionality constant $\frac{1}{\delta}$ and an offset parameter, $B$, representing the basal $OD_{600}$. Now, $f(t)$ is modelled using an exponential function that begins growth after $\tau$ hours and has a growth rate $\beta$,

$$y_t|B,\delta,\sigma,\beta,\tau \sim \text{lognormal}\left(\log\left(B + \frac{f(t)}{\delta}\right), \sigma\right)$$

$$\text{where } f(t) = \begin{cases} f(0) \text{ for } t \leq \tau, \\ f(0)\exp\left\{\beta(t-\tau)\right\} \text{ for } t > \tau. \end{cases}$$

The initial OD at time $t = 0$, $f(0)$, is not a parameter and the true initial fungal inocula are used. All of the parameters are given the same priors as in the Logistic-OD-calibration model (Section B.1)

##### B.2.6 Gompertz-OD-calibration

Again, the OD data, $y_t$, at a time $t$ is modelled as a linear transform of latent fungal growth, $f(t)$ [N/$\mu$l], with proportionality constant $\frac{1}{\delta}$ and an offset parameter $B$. However, $f(t)$ is a now a Gompertz growth model that begins growth after $\tau$ hours and has an initial growth rate $c$,

$$y_t|B,\delta,\sigma,c,K,\tau \sim \text{lognormal}\left(\log\left(B + \frac{f(t)}{\delta}\right), \sigma\right)$$

$$\text{where } f(t) = \begin{cases} f(0) \text{ for } t \leq \tau, \\ f(0)\exp\left\{\log\left(\frac{K}{f(0)}\right)\left(1 - e^{-c(t-\tau)}\right)\right\} \text{ for } t > \tau. \end{cases}$$

where $K$ is the carrying capacity parameter. Again, the initial $OD_{600}$ at time $t = 0$, $f(0)$, is not a parameter and the true initial fungal inocula are used. All of the parameters are given the same priors as in the Logistic-OD-calibration model (Section B.1) and the initial growth rate, $c$, is given the same prior as the growth rate, $\beta$, in the Logistic-OD-calibration model. When $f(0) = 0$ the following model is used

$$y_{t,b}|B,\sigma \sim \text{lognormal}\left(\log(B), \sigma\right)$$

where $y_{t,b}$ is the $OD_{600}$ of the blanks.

##### B.2.7 GP-OD-calibration

As with the Exponential- and Gompertz-OD-calibration models, the $OD_{600}$ data, $y_t$, at a time, $t$, is modelled as a linear transform of latent fungal growth [N/$\mu$l], with proportionality constant $\frac{1}{\delta}$ and a basal $OD_{600}$ of $B$. However, here, $f(t)$, is a GP that models the log of the latent true fungal growth. Hence,

$$y_t|B,\delta,\sigma,\alpha,\rho \sim \text{lognormal}\left(\log\left(B + \frac{e^{f(t)}}{\delta}\right), \sigma\right).$$

As the GP, $\boldsymbol{f}$, is latent in this model, we cannot use the analytical form of the posterior predictive of the derivative of the GP, $\boldsymbol{g}$, at time points $\boldsymbol{t}^d$ as in the GP-OD model. Instead, the derivative of the GP is inferred with the GP, $\boldsymbol{f}$, using the following GP prior [3],

$$\begin{pmatrix} \boldsymbol{f} \\ \boldsymbol{g} \end{pmatrix} \sim \text{MVN}\left(\boldsymbol{0}, \begin{bmatrix} K(\boldsymbol{t},\boldsymbol{t}) & \partial_2 K(\boldsymbol{t},\boldsymbol{t}^d) \\ \partial_1 K(\boldsymbol{t}^d,\boldsymbol{t}) & \partial_1\partial_2 K(\boldsymbol{t}^d,\boldsymbol{t}^d) \end{bmatrix}\right)$$

where $K$ is again the exponentiated quadratic kernel. Finally, for the data from blanks, $y_{t,b}$, we use the following to model the $OD_{600}$,

$$y_{t,b}|B,\sigma \sim \text{lognormal}\left(\log(B), \sigma\right).$$

The same priors as the Logistic-OD-calibration model (Section B.1) are used for $B$, $\delta$ and $\sigma$ and the same prior for $\rho$ is used as in the GP-OD model (Section B.2.4). We used a different prior than the GP-OD model for the output variance, $\alpha$, because the GP in the GP-OD-calibration model represents the log of the latent true fungal growth [N/$\mu$l], which has a different expected range than the GP in the GP-OD model that models the log of the $OD_{600}$ (Section B.2.4). The log of the true fungal growth is expected to vary by at least the same magnitude the initial condition values vary (from $\log(1)$ to $\log(10^3)(= 6.91$ to three significant figures (3.s.f.)) [N/$\mu$l]). We also want the output variance to cover different orders of magnitude as we do not know the true fungal growth. Hence, we used $\alpha \sim \mathcal{N}(5, 2)$ as a prior for $\alpha$.

When fitting the GP-OD-calibration model to only the high initial inoculum $OD_{600}$ data, we ran the Hamiltonian Monte Carlo (HMC) chains in Stan for 5000 iterations with 2000 warm-up iterations, instead of 4000 iterations with 2000 warm-up iterations that was used when the model is fit to all the $OD_{600}$ data, to ensure there were no signs of non-convergence.

###### B.2.8 Mixed-effect logistic-OD-calibration

The model is parameterised the same as the Logistic-OD-calibration model (Section B.1), however we model a background media $OD_{600}$ value, $B^j$, per replicate, $j$,

$$y_t^j | B^j, \delta, \sigma \sim \text{lognormal}\left(\log\left(B^j + \frac{f(t)}{\delta}\right), \sigma\right)$$
$$\text{where } f(t) = \begin{cases} f(0) \text{ for } t \leq \tau, \\ K\left(1 + \left(\frac{K - f(0)}{f(0)}\right) e^{-\beta(t-\tau)}\right)^{-1} & \text{for } t > \tau. \end{cases}$$

The priors for all the parameters are kept the same as in the Logistic-OD-calibration model (Section B.1), apart from the parameters associated with the background media ($B^j$ and $\sigma_B$):

- $B^j$: background $OD_{600}$ value per replicate. We model these as distributed around the sample mean, $\bar{y}_0$ of the blanks, $B^j \sim \mathcal{N}(\bar{y}_0, \sigma_B)$, where $\bar{y}_0 = \frac{1}{N_b}\sum_{i=1}^{N_b} y_i^b$ and $y_i^b$ denotes the $OD_{600}$ values of the blanks and $N_b$ is the total number of data points of the blanks.
- $\sigma_B$: scale of the deviation of the background $OD_{600}$ value per replicate around the sample mean, $\sigma_B \sim \mathcal{N}^+(0, 1)$.

###### B.2.9 No-delay logistic-OD-calibration

The model is parameterised the same as the Logistic-OD-calibration model (Section B.1), however we do not include a delay to the start of logistic growth ($\tau = 0$),

$$y_t | B, \delta, \sigma \sim \text{lognormal}\left(\log\left(B + \frac{f(t)}{\delta}\right), \sigma\right)$$
$$\text{where } f(t) = K\left(1 + \left(\frac{K - f(0)}{f(0)}\right) e^{-\beta t}\right)^{-1}$$

The priors for all the parameters are kept the same as in the Logistic-OD-calibration model (Section B.1).

###### B.2.10 Logistic-HL

We model the hyphal length (HL) data collected from microscopy, $y_{H,t}$, at time $t$ by

$$y_{H,t} | \sigma_H, \beta, K, \tau, f(0) \sim \text{lognormal}\left(\log\left(f(t)\right), \sigma_H\right)$$
$$\text{where } f(t) = \begin{cases} f(0) \text{ for } t \leq \tau, \\ K\left(1 + \left(\frac{K - f(0)}{f(0)}\right) e^{-\beta(t-\tau)}\right)^{-1} & \text{for } t > \tau. \end{cases}$$

where the logistic function has a hyphal growth rate $\beta$, delay $\tau$ and a carrying capacity $K$. The following parameters have the same priors as in the Logistic-OD-calibration model (Section B.1); $\tau$ and $\beta$. The parameters that have different priors are listed below:

- $K$: average well carrying capacity. Since the maximum HL value is around 300$\mu$m, we place a prior on the transform, $L = \log_{10}(K)$, of $L \sim \mathcal{N}^+(4, 2)$. The maximum HL will not be the carrying capacity of the well and we want prior of the carrying capacity to cover different orders of magnitude. Hence, the scale of the prior for this parameter was taken to be 2.
- $\sigma_H$: multiplicative noise scale. We do not expect the noise to exceed $\approx 200\%$ of the of the observed value on average; $\sigma_H \sim \mathcal{N}^+(0, 1)$.
- $f(0)$: the initial HL of a single hypha, which we assume to be close to 0$\mu$m $f(0) \sim \mathcal{N}^+(0, 1)$.

###### B.2.11 Gompertz-HL

We model the HL data, $y_{H,t}$, at time $t$ by

$$y_{H,t}|\sigma_H, c, K, \tau, f(0) \sim \text{lognormal}\left(\log\left(f(t)\right), \sigma_H\right)$$
$$\text{where } f(t) = \begin{cases} f(0) \text{ for } t \leq \tau, \\ f(0)\exp\left\{\log\left(\frac{K}{f(0)}\right)\left(1 - e^{-c(t-\tau)}\right)\right\} \text{ for } t > \tau. \end{cases}$$

where the Gompertz function has an initial growth rate $c$, a delay $\tau$ and a carrying capacity $K$. The carrying capacity, $K$, the initial growth rate, $c$, the delay, $\tau$, the scale of the noise $\sigma_{\text{H}}$, and the parameter for the initial HL, $f(0)$, are given the same priors as the priors of the carrying capacity, $K$, growth rate, $\beta$, the delay, $\tau$, the scale of the noise $\sigma_{\text{H}}$, and the initial HL, $f(0)$, in the Logistic-HL model (Section B.2.10), respectively.

#### B.2.12 GP-HL

We model the log-transformed HL data by a GP: $f(t)$, where $\sigma_H$ is the scale of the observed noise,

$$\{\log y_{H,t}\}\,|\alpha, \rho, \sigma_H \sim \mathcal{N}\left(f(t), \sigma_H\right)$$

The GP is equipped with a zero mean GP prior and the exponentiated quadratic kernel. The hyper-priors for $\rho$ and $\sigma_H$ are the same as for $\rho$ and $\sigma$ in the GP-OD model (Section B.2.4), respectively. For $\alpha$, the output scale of the GP, we use $\alpha \sim \mathcal{N}^+(0, 1)$ because the HL data varies on a larger scale than the $OD_{600}$. Finally, the derivative of the fitted GP and growth rate are calculated as described for the GP-OD model.

###### B.2.13 Logistic-NC

We model the nuclear count (NC) data, $y_{N,t}$, at time $t$ by

$$y_{N,t}|\sigma_N, \beta, K, \tau, f(0) \sim \text{lognormal}\left(\log\left(f(t)\right), \sigma_N\right)$$
$$\text{where } f(t) = \begin{cases} f(0) \text{ for } t \leq \tau, \\ K\left(1 + \left(\frac{K - f(0)}{f(0)}\right)e^{-\beta(t-\tau)}\right)^{-1} \text{ for } t > \tau. \end{cases}$$

where the logistic function has a growth rate $\beta$, a delay $\tau$, and a carrying capacity $K$. The following parameters have the same priors as in the logistic-OD-calibration model (Section B.1); $\tau$ and $\beta$. The parameters that have different priors are listed below:

- $K$: average well carrying capacity. A prior is placed on a transform, $L = \log_{10}(K)$, $L \sim \mathcal{N}^+(2, 2)$. This is because we want the carrying capacity to be higher than the maximum observed nuclear count ($\approx$30) but also cover different orders of magnitude.
- $\sigma_N$: multiplicative noise scale. We do not expect the noise to exceed $\approx$ 200% of the of the observed value on average; $\sigma_N \sim \mathcal{N}^+(0, 1)$.
- $f(0)$: the initial NC per hypha, which we assume to be close to 0, $f(0) \sim \mathcal{N}^+(0, 1)$.

###### B.2.14 Gompertz-NC

We model the NC data, $y_{N,t}$, at time $t$ by

$$y_{N,t}|\sigma_N, c, K, \tau, f(0) \sim \text{lognormal}\left(\log\left(f(t)\right), \sigma_N\right)$$
$$\text{where } f(t) = \begin{cases} f(0) \text{ for } t \leq \tau, \\ f(0)\exp\left\{\log\left(\frac{K}{f(0)}\right)\left(1 - e^{-c(t-\tau)}\right)\right\} \text{ for } t > \tau. \end{cases}$$

where the Gompertz function has an initial growth rate $c$, a delay $\tau$ and a carrying capacity $K$. The carrying capacity, $K$, the initial growth rate, $c$, the delay, $\tau$, the scale of the noise $\sigma_{\text{N}}$, and the parameter for the initial NC, $f(0)$, are given the same priors as the priors of the carrying capacity, $K$, growth rate, $\beta$, the delay, $\tau$, the scale of the noise $\sigma_{\text{N}}$, and the initial NC, $f(0)$, in the Logistic-NC model (Section B.2.13), respectively.

### B.2.15 GP-NC

We model the log-transformed NC data using a GP $f(t)$, where $\sigma_N$ is the scale of the observed noise,

$$\log y_{N,t}|\alpha, \rho, \sigma_N \sim \mathcal{N}\left(f(t), \sigma_N\right)$$

We, again, use a zero mean GP prior and the exponentiated quadratic kernel. In addition, the derivative of the GP and growth rate are calculated as described for the GP-OD model (Section B.2.4). The hyperpriors for $\alpha$ and $\rho$ are the same as in the GP-OD model. For $\sigma_N$, the scale of the measurement noise we use $\sigma_N \sim \mathcal{N}^+(0,1)$ as we expect the NC data to have larger scale of experimental noise than the $OD_{600}$ data.

### C Random walk model comparison

We also compare all reference models with a random walk model (as a sanity check), where the random walk for the observed $OD_{600}$, $y_t^j$, is specified as $y_t^j \sim \mathcal{N}(y_{t-1}^j, \sigma)$, for the $j$-th replicate at time $t$, and $\sigma$ is the scale of the measurement noise to be inferred. The prior for $\sigma$ is kept the same as the prior used in our Logistic-OD-calibration model (Section B.1). Our model (Section B.1) and the considered reference models (Section B.2) have lower (better) RMSE and relative LPD than a random walk model that has median testing $\log_{10}$ RMSE and $\log_{10}$ relative LPD values of 0.327 and 2.73, respectively (both to 3.s.f.).
